# Combined climate change and arsenic pollution increases rice cadmium with little effect on micronutrients

**DOI:** 10.64898/2026.10.28.762300

**Authors:** Jill Bachelder, Tianmei Wang, Scott Fendorf, E. Marie Muehe

**Author notes:** Corresponding author Marie Muehe, Plant Biogeochemistry, Department of Applied Microbial Ecology, Helmholtz Centre for Environmental Research UFZ, Permoserstr. 15, 04318 Leipzig, Germany.

## Abstract

**(Background and Aims):** Rice production faces the dual threat of increasing soil As contamination and climate change. Each stressor can alter micronutrient (e.g. Fe, Zn) and toxin (e.g., Cd) plant uptake and internal translocation. Yet, the combined effects of soil As contamination and climate change factors on the nutritional quality and safety of rice grain remain poorly elucidated.

**(Methods):** A full-factorial pot experiment growing rice under two soil As levels (low-As, high-As) and four climate conditions (today; elevated CO_2_ (+435 ppm CO_2_); elevated temperature (+5°C); future (combined elevated CO_2_ and temperature)) was performed.

**(Results):** Grain Fe decreased slightly with soil As contamination and was barely altered by future climate conditions. Grain Zn and Cd both decreased with soil As contamination and increased with future climate. For grain Zn, the decrease with As contamination led to an overall decrease (by 25%) with combined soil As contamination and future climate. For Cd, increased concentrations under future climate outweighed the effect of soil As, leading to two-fold increased grain concentrations with As contamination and future climate combined.

**(Conclusion):** Our results provide insight into the processes controlling Fe, Zn, and Cd in rice grown in polluted systems threatened by climate change, and highlight the critical importance of multifactorial, mechanistic studies.

## Introduction

Rice is a staple crop and essential source of protein and nutrients for many people worldwide. However, rice contains low concentrations of essential micronutrients such as Fe and Zn compared to human dietary requirements (Liu et al. 2024), and often problematic concentrations of toxic, non-essential metals such as As and Cd (Zhao and Wang 2020). The balance between maintaining adequate micronutrient concentrations and limiting toxin is a major challenge in rice production, as micronutrient and toxin concentrations can be affected by soil contamination and climate change (Williams et al. 2007; Zhu et al. 2018; Luo et al. 2019; Wang et al. 2020; Zhou et al. 2023).

In rice production systems, soil can be contaminated by geogenic As and anthropogenic activity, e.g. via irrigation with As-contaminated water and pesticide use (Meharg and Rahman 2003; Islam et al. 2004; Hang et al. 2009; Dittmar et al. 2010; Neumann et al. 2011; Muehe and Kappler 2014; Limmer and Seyfferth 2024; Zhang et al. 2024). Paddy soil As contamination can lead to reduced rice yields due to its toxicity (Panaullah et al. 2009). Arsenic contamination has been shown to decrease micronutrient concentrations in rice (Williams et al. 2009). Environmental factors associated with climate change can also affect rice grain yields and nutritional quality. For example, elevated CO_2_ may dilute micronutrient concentrations by increasing biomass production (Myers et al. 2014; Zhu et al. 2018; Yang et al. 2024), while elevated temperature may concentrate micronutrients by decreasing grain biomass (Wang et al. 2020; Bi et al. 2024). Though the threats of soil contamination and climate change are well understood individually, few studies have investigated their combined effects on rice grain yield and nutrient quality and overall grain safety. Combined As stress and climate change conditions nearly doubled inorganic As concentrations in grains compared to low-As, ambient climate controls (Muehe et al. 2019; Farhat et al. 2021; Wang et al. 2025; Duncan et al. 2025). Yet, these studies did not assess other key determinants of grain quality such as micronutrients (Fe, Zn) or additional toxins (e.g. Cd). Thus, a multifactorial climate study investigating the combined effects of climate change and soil As contamination on both micronutrient and toxic metal accumulation in rice is still lacking.

Arsenic contamination and future climatic conditions have potential to alter the processes controlling uptake of micronutrients and toxins in rice. There is growing evidence that climate change can transform the speciation of metals such as Fe, Zn, and Cd to forms readily taken up by plants (i.e., made phytoavailable) (Pieńkowska et al. 2025; Wang et al. 2026). For example, porewater Cd increased in oxic agricultural soils under combined elevated CO_2_ and temperature compared to today’s climate (Drabesch et al. 2024, 2026). Decreased soil pH and increased dissolved organic carbon promoted Cd desorption and soluble complexes. However, these findings are limited to oxic soil systems, where Cd is primarily associated with clays, iron (hydr)oxides, or carbonates. In contrast, water-logged soils typical of rice production are governed by reducing conditions, under which reductive dissolution of iron (hydr)oxides releases Fe²⁺ and associated elements such as As into porewater (Muehe et al. 2019), especially under further climatic conditions. Simultaneously, Zn and Cd tend to form metal sulfides (ZnS, CdS) of limited solubility and become less phytoavailable (Khaokaew et al. 2012; Fulda et al. 2013; Yin et al. 2025). Whether climate-amplified increases in porewater As and Fe affect the phytoavailability of micronutrients like Zn and Cd in flooded rice-production systems is unknown but crucial for assessing rice grain quality.

Combined climate change and As stress may also affect internal metal translocation in rice. Micronutrients like Fe and Zn are taken up and translocated via designated transporters (Zhao and Wang 2020). In contrast, Cd can exploit multiple micronutrients transporter pathways (e.g., for Fe, Zn, Mn) and move passively with water flow (Hart et al. 2002; Nakanishi et al. 2006; Zhao and Wang 2020). This often leads to higher Cd accumulation relative to essential nutrients like Fe and Zn (Simmons et al. 2003; Zhao and Wang 2020). Arsenic may disrupt transport of micronutrients and Cd via secondary effects due to its general toxicity to plant health which disrupts metabolic pathways (Shaibur et al. 2016). Climate factors such as elevated temperature can intensify Cd translocation to grains (Ge et al. 2019; Zhou et al. 2023). However, no study has assessed the combined effects of climate factors and As contamination on micronutrient (Zn, Fe) and toxic metal (Cd) translocation in rice.

This study aimed to determine how combined climate change and soil As stress affect processes controlling transfer of Fe, Zn, and Cd from soil to rice grains. Rice plants were cultivated under future climatic conditions (combined elevated CO_2_ +435 ppm and elevated temperature +5°C) in both native and As-spiked soil. To expand our mechanistic understanding of metal movement from soil to grain, additional treatments with elevated CO_2_ or elevated temperature alone were included. To link potential elemental phytoavailability to grain contents, metal concentrations in soil porewater were measured throughout the experiment. At maturity, metal partitioning in plant organs (i.e., straw, panicles, husks, grains) were quantified to evaluate translocation. These results offer crucial insight into the combined potential effects of soil As and climate change on the entry of micronutrients and toxic elements into food systems via rice.

## Materials and Methods

### Study design

This study expands on the experiment conducted by Muehe et al. (2019), which examined the effects of future climatic conditions on rice yield and grain As content. In brief, the experiment used a clay paddy soil from Arbuckle, California, US (39°01’38.2"N 121°55’36.7"W, elevation: 22 m), cultivated with the commonly used California rice variety *Oryza sativa* L. sub-species japonica, cultivar Calrose, variety M206. The soil’s characteristics included a pH of approximately 6.9, a cation exchange capacity of 76 cMolc kg^-1^, and total carbon and nitrogen contents of 13 g kg^-1^ and 1 g kg^-1^, respectively (see Table S1 and Muehe et al. (2019)). The native As content of the paddy soil was 7.3 mg kg^-1^ . To assess the combined effects of climate change condition and soil As on plant performance, the soil was spiked to a concentration of 24.6 mg As kg^-1^ using 80% arsenite (NaAsO, analytical grade) and 20% arsenate (Na_2_HAsO_4_x7H_2_O, analytical grade), followed by a four months acclimatization period under each climatic condition (see next paragraph). Soil metal concentrations were determined using X-ray fluorescence on 5 g of freeze-dried and powdered soil samples with a Spectro XePositive HE XRF Spectrometer (AMETEK, Germany) and are presented in Table S1.

Rice plants were cultivated under controlled environmental conditions within growth chambers (1.2 × 1.8 × 1.8 m, polycarbonate) housed inside two larger greenhouses. Each greenhouse contained four growth chambers. Chamber 1 simulated today’s climate based on conditions of Californian rice-growing regions in 2016 (33 °C average high daytime temperature during rice-growing season from May to September) and atmospheric CO_2_ of 415 ppm_v_). Chamber 2 simulated future climatic conditions according to the IPCC RCP 8.5 scenario (38 °C and atmospheric CO_2_ of 850 ppm_v_), which was projected as the most realistic scenario in the 2021 report (Friedlingstein et al. 2022). In chambers 3 and 4, either CO_2_ or temperature was elevated individually. Further details on the climate control methods for both atmosphere and soil are described in Muehe et al.(2019).

Approximately 3.4 kg of soil was placed into high-density polyethylene pots (14-cm diameter, 18-cm height) filled to 4 cm below the rim. The soils were kept water-logged to the rim using pH 7 tap water. The M206 rice variety was germinated in sterilized tap water under the respective climatic conditions. After one week, three seedlings with similar shoot and root lengths were transplanted into each pot, with eight pots allocated per environmental condition. Urea (CON_2_H_4_, analytical grade) was applied as fertilizer at a rate of 185 kg nitrogen ha^−1^ at the start of the season, after 4.5 weeks of growth, and at panicle initiation, mirroring the practices of Californian rice farmers. To prevent biases in light exposure, temperature, and atmospheric CO_2_ due to position of the plants within and across the chambers, pots were rotated weekly within the basins, and the chambers themselves were alternated every 3–4 weeks.

### Porewater sampling and analysis

Porewater was extracted using Rhizon samplers (0.15 µm pore size, RhizonFlex 19.21.25, Rhizosphere Research Products), which were inserted horizontally at the midpoint of each pot. The extracted porewater was collected in acid-washed, butyl-stoppered, evacuated glass vials. In an anoxic chamber (96% N_2_/4% H_2_ atmosphere), porewater was filtered through 0.2-µm syringe filters (polyethersulfone membrane, Minisart® High Flow, Sartorius, US) and diluted in 2% nitric acid (analytical grade) for total elemental quantification with inductively coupled plasma optical emission spectroscopy (ICP-OES, iCAP6000, Thermo Scientific, UK). Details and data on other porewater parameters such as As speciation, pH, DOC, DIC can be found in (Muehe et al. 2019).

### Plant harvest and analysis

At harvest, rice plants were cut 2 cm above the water level, wrapped in butcher paper, and dried at 40 °C in the dark for four weeks. The dried plant material was separated into grains, panicles and straw (which included combined leaves and stems), and each fraction was weighed. Husks were removed from the grains by rubbing filled spikelets between two plastic crusher plates. Grains, husks, and straw were ground using ball and cutting mills followed by liquid nitrogen grinding if necessary. For elemental analysis, between 0.05 and 0.5 g of each plant tissue type was digested in a microwave digester (Mars 6 Digester, CEM, US) using 3 mL of 65 % nitric acid (HNO_3_, analytical grade) and 2 mL of 30 % H_2_O_2_ (analytical grade). The digestion process included a 15-min ramp phase to 180 °C, followed by a 15-min hold at this temperature, and a 30 min cool-down phase. The digested samples were diluted with double-deionised water, and elemental concentrations were measured using inductively coupled plasma mass spectrometry (single quadrupole, XSeries2, Thermo Fisher Scientific Inc., USA). The European Reference plant material ERM-BC211 was included for extraction efficiency checks; and for instrumental stability and elemental recovery checks regular QCs in the mid-calibration range and additional 2 ppb As spikes were added. Total elemental contents per tissue (straw, panicle, husks, and grain) were calculated by multiplying the elemental grain concentration by the grain yield, i.e. following equation 1:

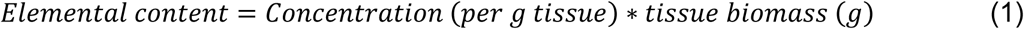

The straw retention fraction for each element was calculated following equation 2:

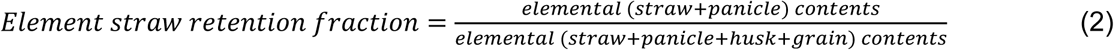

The husk retention fraction for each element was calculated following equation 3:

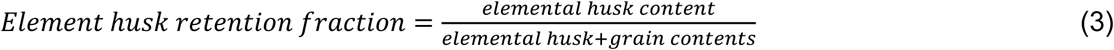

Panicle elemental retention fractions were not calculated.

### Statistics

Data were analyzed using the R programming language (version 4.5.2) and RStudio (version 2026.01.0+392). Mean values and standard deviations were calculated from eight biological replicates, except for grain Cd where only four replicates were included (Table S2). Boxplots were generated using ggplot2 to display the median, interquartile range (IQR), and whiskers extending to the most extreme values within 1.5 × IQR. Statistical analysis was performed using a generalized least squares (GLS) model at a confidence level of 0.95. For samples whose residuals were shown by Levene tests to be of non-homogenous variance, a constant variance function (varIdent) was employed to eliminate the effect of heteroscedasticity from the low-vs. high-As treatments. To determine whether the two variables (As and climate condition) and their interactions had a significant effect on the data, an ANOVA type II was performed using the GLS output. An emmeans post-hoc test was used to compare soil As levels and climatic conditions at a confidence level of 0.95. A summary of all ANOVA and emmeans posthoc outputs per response variable is provided in Table S3.

Principal component analysis (PCA) was used to characterize elemental partitioning and biomass allocation within rice tissues, considering the elements Fe, Zn, and Cd. Calculations were performed using R (version 4.4.1). The following seven variables were included to explain the variance in grain elemental concentration: straw, panicle, and husk elemental concentrations, and corresponding straw, panicle, husk, and grain biomass. All variables were standardized to unit variance prior to PCA. PCA was performed using the prcomp function on the scaled data. Principal component (PC) scores were extracted and their structure was evaluated using (i) variance explained, (ii) squared loadings to quantify tissue contributions, and (iii) bootstrap resampling to assess loading stability. Bootstrapping (n = 1000) was performed by sampling observations with replacement, repeating PCA on each resample, and aligning resulting loadings to the reference PCA using sign correction based on correlation with reference loadings. Mean loadings and 95% bootstrap confidence intervals were calculated across re-sampled datasets. Loadings were considered stable when confidence intervals did not substantially overlap zero, with loading values within ±0.05 treated as negligible to avoid over-interpreting trivial sign fluctuations. To evaluate biological relevance of partitioning axes, linear mixed-effects models (lmm) were used to test associations between PC scores and grain elemental concentrations, including climate condition as a fixed factor. Combined models were fitted for relevant partitioning effects alone (PC1–PC4 + climate, when ANOVA positive) and exposure (porewater elemental area under the curve). Model inference was based on parameter estimates, ANOVA (Type III tests), estimated marginal means for climate condition effects, and marginal R² calculated from observed versus fitted values.

## Results

### Grain quality

Grain biomass, published previously in Muehe et al. (2019), was approximately 40% less when produced under high compared to low-As (Table S4). Climate conditions had little effect on grain biomass when compared within As treatments, but the combined effect of high-As and elevated temperature (38°C) or future conditions (38°C, 850 ppmv CO_2_) led to significant decreases in grain biomass production of approximately 42-65% compared to low-As under today (33°C, 415 ppmv CO_2_), and was significant for interaction according to the ANOVA test.

Grain Fe concentrations averaged 19±2 mg Fe kg^-1^ under low-As and today (Fig. 1a), with no significant difference between today and any other climatic. In the high-As soil with today’s climate, grain Fe averaged 17±1 mg Fe kg^-1^ . In the high-As soil with elevated temperature, grain Fe reached 20±2 mg Fe kg^-1^, which was statistically higher than today’s climate under high-As. In both low-and high-As soils, grain Fe was significantly lower with elevated CO_2_ compared to both elevated temperature and future conditions.

**Fig. 1:**
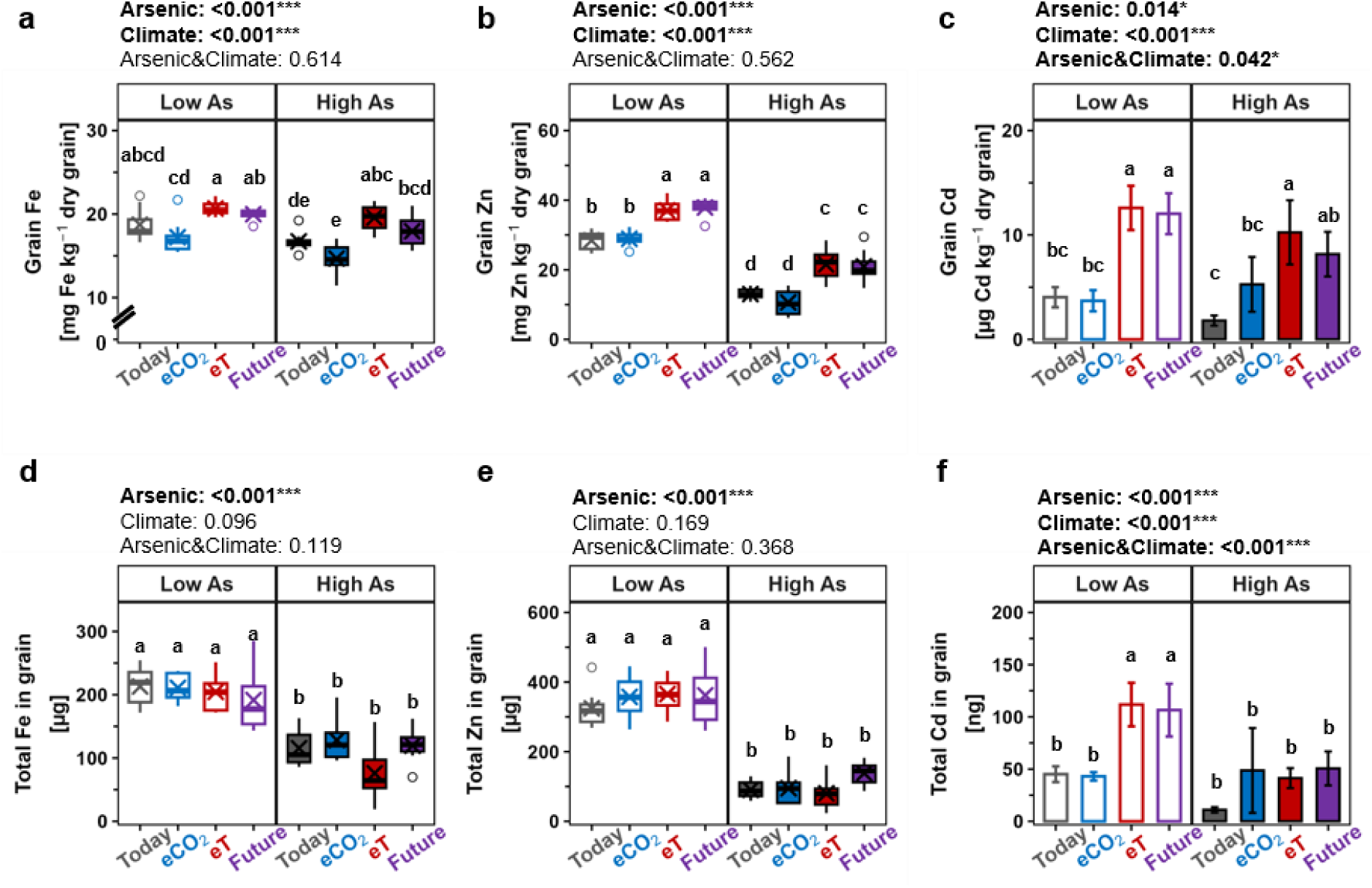
Elemental concentrations in Oryza sativa L. cv. M206 grains produced under varying climatic and soil arsenic conditions. Concentrations of (a) iron, (b) zinc, and (c) cadmium in dehusked grains are presented. Total grain contents of (d) iron, (e) zinc, and (f) cadmium are also provided. Rice was subjected to today’s (33°C, 415 ppmv CO2; grey), elevated CO2 (33°C, 850 ppmv CO2; blue), elevated temperature (38°C, 415 ppmv CO2; red), and future (38°C, 850 ppmv CO2; purple) climatic conditions with low (no fill, 7.3 mg As kg^−1^) and high (filled, amended to 24.5 mg As kg^−1^) soil arsenic levels. Iron and zinc included eight biologicalreplicates, cadmium included four. Boxplots show the median, interquartile range (IQR), and whiskers extending to the most extreme values within 1.5 × IQR; observations outside this range are plotted as outliers (circles). Barplots show the mean ± standard deviation. Statistical analyses of climate, arsenic, and their interaction on elemental grain concentrations and contents were performed using ANOVA at a 95% confidence level using the results of a generalized least squares test. Results are shown above each plot, with significant findings highlighted in bold. Post-hoc comparisons using the Emmeans test are indicated by different lowercase letters, where letters denote statistically significant differences (p<0.05). Details of analytical replicates and statistical calculations are provided in Tables S2 and S3, respectively.

Grain Zn concentrations averaged 28±3 mg Zn kg^-1^ under low-As at today’s climate, (Fig. 1b). Low-As combined with elevated temperature or future conditions increased grain Zn concentrations by approximately 35% to approximately 38 mg Zn kg^-1^ compared to low-As and today’s climate. In the high-As soil with today’s conditions, grain Zn averaged 13±2 mg Zn kg^-1^ . This was significantly lower compared to grain Zn under low-As and today’s climate, constituting a decrease of 54%. High-As and future conditions led to 21±5 mg Zn kg^-1^, an increase of 62% compared to high-As and today’s conditions, but combined high-As and future condition, a 25% decrease in grain Zn concentrations was observed compared to low-As and today’s climate. In both low-and high-As soils, elevated CO_2_ tended to have no effect on grain Zn concentrations compared to today’s climate, while elevated temperature led to similar Zn grain concentrations as future climatic conditions. Thus, the increased Zn concentrations triggered by future condition were outweighed by high-As.

Grain Cd averaged 4±1 µg Cd kg^-1^_dry grain_ in the low-As soil and today’s climatic conditions (Fig. 1c). Future climatic conditions increased grain Cd by three-fold compared to today’s climate in the low-As soil, i.e. to 12±2 µg Cd kg^-1^_dry grain_. With today’s climate, grain Cd decreased in high-compared to low-As soil by 55%, to a mean of 1.8±0.5 µg Cd kg^-1^_dry grain_. The ANOVA test indicated a significant effect of As contamination on grain Cd, though pairwise comparison did not show a significant difference. In the high-As soil, grain Cd concentrations increased four-fold with future condition compared to today’s climate, to a mean value of 8±2 µg Cd kg^-1^ . Future conditions led to a much larger increase in grain Cd compared to the increase observed for grain Zn (i.e., >three-fold increase in Cd compared to ∼30-60% increase in Zn) compared to today’s conditions within each As treatment. Hence, even though grain Cd concentrations followed similar overall trends as grain Zn, the effect of climate condition was much higher on Cd than Zn, as also evidenced by a significant As-climate interaction in the ANOVA for grain Cd (Fig. 1c). Combined high-As and future climatic conditions led to a two-fold increase in grain Cd concentrations compared to low-As and today’s climate, though this difference was not statistically significant given data variance of only four usable replicates in the dataset.

For Fe and Zn, expressing grain elemental accumulation as total grain content (i.e., concentrations multiplied by biomass) removed all significant effects of climatic conditions (Fig. 1d,e). Across all climatic conditions, Fe content averaged 205±35 µg Fe kg^-1^ in the low-As soil. Soil As concentrations, by contrast, had a pronounced impact on Fe, which decreased by 46% to 110±39 µg Fe kg^-1^ in the high-As soil. The decrease in grain Zn content with As contamination was even more extreme, with grain Zn content decreasing from 352±65 µg Zn kg^-1^ in the low-As soil across all climate conditions to 101±44 µg Zn kg^-1^ in the high-As soil (71% decrease).

Unlike Fe and Zn, expressing grain Cd accumulation as total grain content did not remove the effect of climate (Fig. 1f). With low-As and today’s climate, grain Cd content averaged 45±8 ng Cd kg^-1^, with similar findings under elevated CO_2_. This increased 2.3-fold to 106±25 ng Cd kg^-1^ in the low-As soil and future condition, which was primarily driven by elevated temperature. ANOVA identified significant effects of both soil As and climate on grain Cd content, as well as a significant interaction between the two factors (Fig. 1f). However, these effects were not consistently reflected in pairwise comparisons. For example, no significant difference in grain Cd content was detected between low-and high-As soils under today’s climate, and the interaction effect was not clearly supported by pairwise comparisons within the high-As soil. This discrepancy likely reflects the lower replication of the grain Cd dataset (n = 4) compared with Fe and Zn (n = 8). Nevertheless, grain Cd content in the high-As soil showed a tendency to increase under future conditions compared to today’s climate.

Both concentrations and contents of Zn decreased in all aboveground organs grown in high-As compared to low-As soils (Fig.s S1-S2). There was no effect of As contamination or climate on Fe and Cd concentrations and contents in husks, panicles, and straw (Fig.s S1-S2).

### Soil porewater

In the low-As soil with today’s climate, porewater Fe concentrations increased incrementally until week 11, reaching a concentration of 11±1 mg L^-1^, which remained stable until the end of the growth period (Fig. 2a). Low-As and elevated CO_2_ had similar effects on porewater Fe but stabilized 4 weeks earlier than under today’s climate. In the low-As soil with elevated temperature, porewater Fe peaked during the early stages of the growing period, reaching 32 ± 5 mg L^-1^ at week 4. Porewater Fe in the low-As soil with future climate conditions followed similar trends, though peak values were 28% lower in comparison. After peaking in both high-temperature conditions, porewater Fe decreased by week 9 with half the porewater Fe concentration as today’s climate. Comparing low-and high-As soils, high-As and today’s climate resulted in a faster increase in porewater Fe concentrations compared to low-As and today’s climate and remained stable at approximately 13 mg L^-1^ from week 4 onward. Generally, climate condition barely impacted porewater Fe in the high-As soil compared to today, with only a slight increase in porewater Fe at week 4 for the future condition. The trends in porewater Fe were reflected in concentrations of porewater P (Fig. S3a) and trends also generally confirmed by redox active Mn. In the low-As soil, peak porewater Mn concentration was higher and shifted to earlier time points with both high-temperature conditions, and returned to almost no porewater Mn by week 10. In the high-As soil, porewater Mn was greater under all climate conditions and took longer to decrease. Porewater Mn increased early in the growth period and peaked between weeks 3-4 in all treatments regardless of As or climate conditions (Fig. S3d). All climate conditions increased porewater Mn in the low-As soil compared to today’s climate, a dynamic not observed in the high-As soil.

**Fig. 2:**
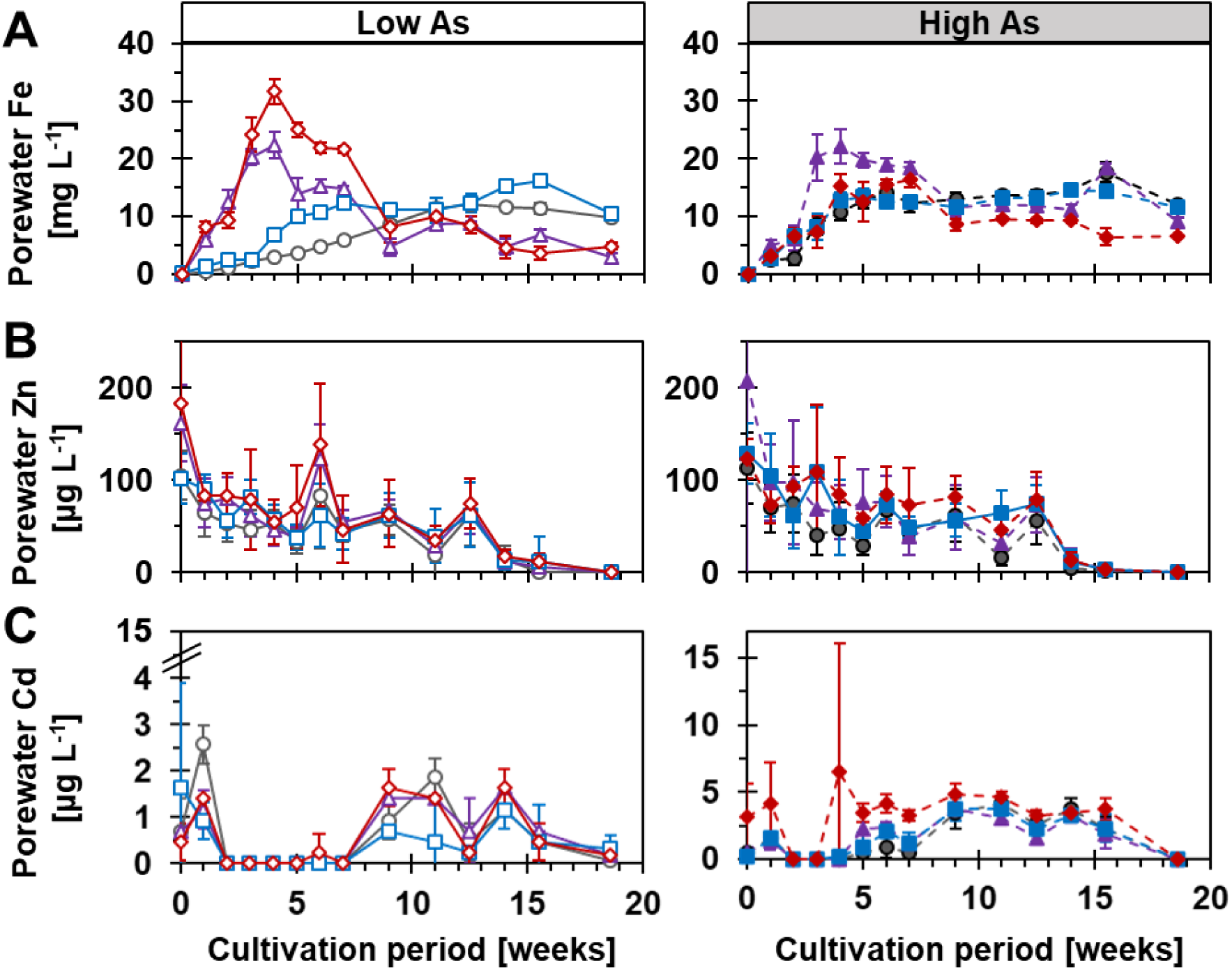
Time-resolved dissolved elements in rhizosphere porewater of Oryza sativa L. cv. M206 grown under varying climatic and soil arsenic conditions. Total porewater (a) iron, (b) zinc, and (c) cadmium extracted from the rhizosphere of rice plants at 10-cm depth using are presented. Rice was subjected to today’s (33°C, 415 ppmv CO2; grey), elevated CO2 (33°C, 850 ppmv CO2; blue), elevated temperature (38°C, 415 ppmv CO2; red), and future (38°C, 850 ppmv CO2; purple) climatic conditions with low (left, 7.3 mg As kg^−1^) and high (right, amended to 24.5 mg As kg^−1^) soil arsenic levels. At each sampling point, the mean ± standard deviation of six biological replicates was calculated. The Fe dataset is published in the supplement of Muehe et al. (2019).

Porewater Zn followed a different trajectory than Fe (Fig. 2b). The initial 100-200 μg Zn L^-1^ decreased to zero by the end of the growth period. Neither soil As nor climate condition altered porewater Zn compared to today’s climate. This followed a similar trend as porewater S (Fig. S3c), which was also highest at the beginning of the cultivation period and gradually decreased to zero by week 10. Soil As did not affect porewater S, but high-temperature conditions deceased porewater S faster compared to both low-temperature treatments.

Similar to Zn but contrary to Fe, porewater Cd was not greatly altered by high-As or future climate condition compared to low-As and today’s climate (Fig. 2c). Under all soil As and climate condition, porewater Cd remained below 6 μg L^-1^. In both low-and high-As soils, porewater Cd remained low until weeks 5-7 and then increased during the second half of the cultivation period, followed by a decrease to zero at the end of the experiment. Given Zn’s ability to bind to clay minerals, porewater K was considered. Porewater K followed a similar trend, i.e. lower concentrations at the beginning of the growing period and high values at the end with high-As leading to an earlier increase than in the low-As soil (Fig. S3b).

### Principal component analysis

To investigate the relative importance of factors associated with grain Fe, Zn, and Cd concentrations, PCA was performed using biomass and elemental concentration data from straw, panicle, and husks (Fig. 3). The principal component (PC) loadings and their bootstrap stabilities are provided in Fig. S4 and Table S5. For all elements, PC1 was dominated by biomass variables (i.e., masses of panicles, straw, husks, and grains). The only notable exception was Zn produced in the high-As soil, for which husk Zn concentration (11%) contributed moderately to PC1. For Fe and Zn, PC1 explained 40.7-45.0% of total data variance, while for Cd it explained 56.8-61.6% of data variance in both low-and high-As soils. Across all elements in the low-and high-As soils, PC2 explained 21.8-28.7% and PC3 explained 8-3-17.6% of data variance (Fig. 3 and S5). Both PCs were primarily associated with elemental concentrations in husks, panicles, and straw, with only sporadic and minor contributions from a biomass variable. The specific tissue concentrations contributing to PC2 or PC3 varied among elements and soil treatments, reflecting the variable bootstrap stability of the loadings (Fig. S4). Owing to the variable bootstrap stability of individual loadings, we adopted a conservative biological interpretation, whereby PC1 was considered to represent biomass-related variation, while PC2 and PC3 were collectively considered to represent variation in tissue elemental concentrations.

**Fig. 3:**
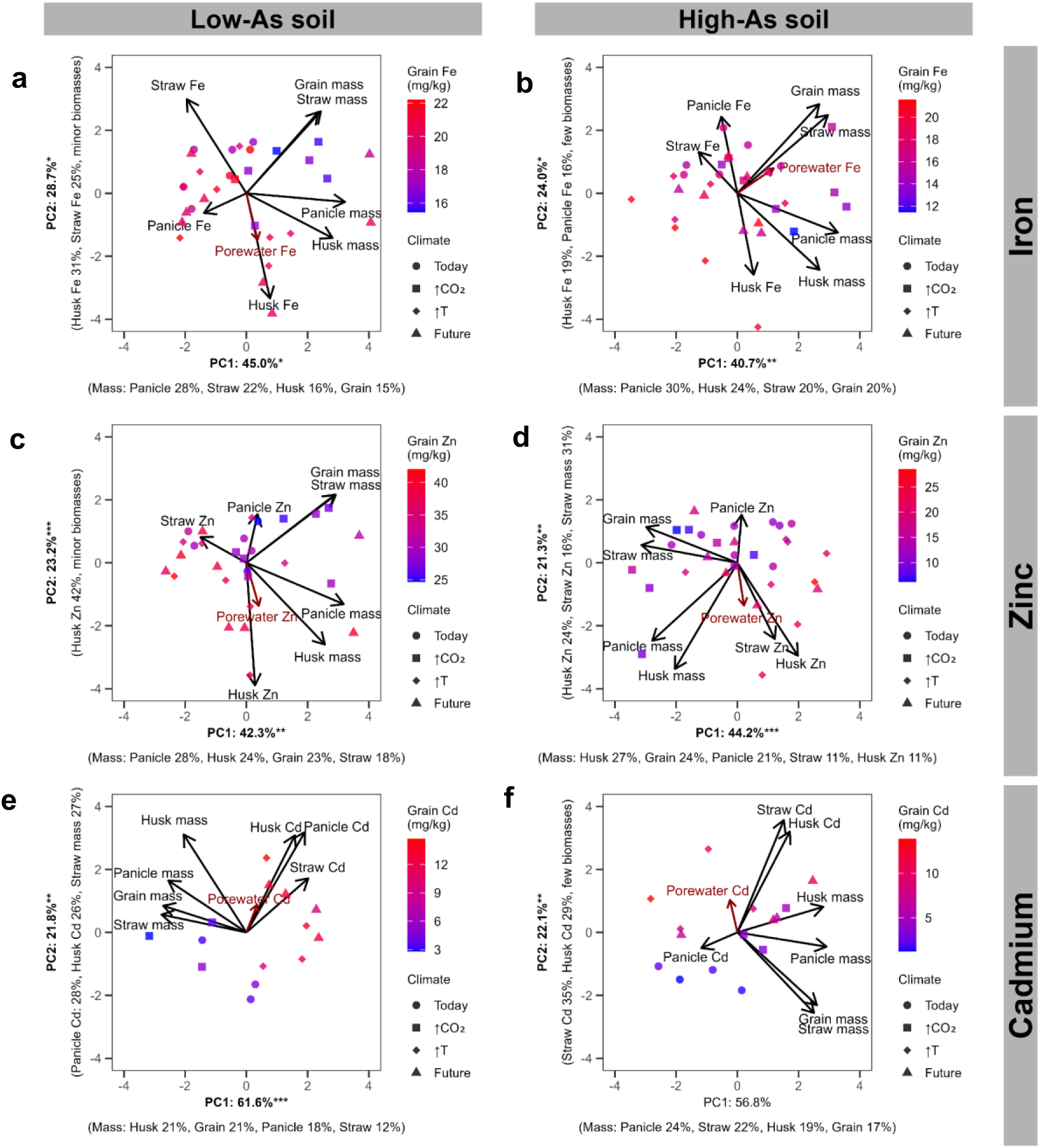
Elemental partitioning structure of rice grown on As-bearing paddy soils under climate change conditions. Principal component analyses (PCA) of (a,b) iron, (c,d) zinc, and (e,f) cadmium partitioning among rice tissues grown in low-As (left panels) and high-As (right panels) soils under today’s (circle, 33°C, 415 ppmv CO2), elevated CO₂ (↑CO₂, squares, 33°C, 850 ppmv CO2), elevated temperature (↑T, diamond, 38°C, 415 ppmv CO2;), and combined future (triangle, 38°C, 850 ppmv CO2;) climate conditions. PCA was performed on scaled tissue element concentrations (straw, panicle and husk) together with biomass allocation traits (straw, panicle, husk and grain mass) to characterize whole-plant elemental distribution independent of absolute magnitude. Points represent individual experimental replicates, colored by grain element concentration and shaped by climate condition. Arrows denote variable loadings in PC1-PC2 space indicating the direction and strength of tissue contributions to multivariate partitioning axes. The red vector represents porewater elemental exposure projected onto PCA space, illustrating the alignment between environmental availability and plant partitioning strategies. Axis labels indicate the percentage of explained variance and dominant tissue contributors. Bold axis labels with significance symbols denote PCs that significantly explained variation in grain element concentration in generalized least-squares models (*p < 0.05, **p < 0.01, ***p < 0.001) (Table S5). Together, the first two PCs capture dominant trade-offs between biomass 15 allocation and tissue elemental enrichment across As and climate conditions. Loading stability according to bootstrapping and partitioning along PC2 and PC3 are presented in Fig.s S4 and S5.

To evaluate whether PCs, climate condition, and soil availability explained grain concentrations, ANOVA/GLS was performed. A significant correlation was found between PC1 (i.e., biomass) for grain Fe and Zn concentrations in both high-and low-As soils, while grain Cd only correlated significantly with PC1 in the low-As soil but not in the high-As soil. For all elements under both low-and high-As, a significant correlation was found between PC2 (i.e., elemental concentrations with sporadic influence of biomass) and grain concentrations. For grain Fe and Zn in the low-As soil, significant correlations were also found with PC3 (i.e., driven by elemental concentrations with no influence of biomass; Fig. S5). No significant correlations were found between PC3 and grain Cd in the low or high-As soils, nor Fe and Zn in high-As. When a PC indicated a significant contribution to grain element, we investigated with ANOVA whether climate affected that PC (Table S5), allowing us to indirectly elucidate a climate impact on grain element through the biological meaning of the respective PC. For both grain Fe and Zn, climate significantly affected PCs related to tissue elemental concentrations in low-As soils, and PC1 related to biomasses in high-As soils. For Cd, climate significantly affected PC1 (i.e. biomass) in low-As soils and PC2 (i.e. element concentration) in high-As soils.

For grain Fe in low-As soil, PC1–PC3 (representing biomass and tissue elemental concentrations) were significant predictors, whereas porewater Fe and climate condition were not (Table S5A). When assessing the effect of climate condition on the significant PCs, only PC2 (primarily representing tissue elemental concentrations, with minor contributions from biomass) was significantly affected by climate condition. For grain Fe in high-As soil, PC1, PC2, and climate condition were significant predictors, whereas porewater Fe was not (Table S5B). The climate effect was driven mainly by the pronounced difference between the elevated CO₂ and future-climate treatments. Both PCs were also directly significantly affected by climate condition.

For grain Zn in low-As soil, PC1–PC4 and climate condition were significant predictors, whereas porewater Zn was not (Table S5C). The climate effect was attributable to differences among temperature regimes. Of the four PCs, only PC3 (primarily representing tissue Zn concentrations) was significantly affected by climate condition. In high-As soil, grain Zn was significantly explained by PC1, PC2, climate condition, and porewater Zn (Table S5D). Here, the climate effect reflected differences between the elevated CO₂ treatment and both high-temperature treatments. However, only PC1 was directly affected by climate condition.

For grain Cd in low-As soil, PC1 and PC2 were significant predictors, whereas climate condition and porewater Cd were not (Table S5E). Nevertheless, climate condition significantly affected PC1. In high-As soil, grain Cd was significantly explained only by PC2, representing variation in tissue Cd concentrations, whereas PC1, climate condition, and porewater Cd were not significant (Table S5F). Although climate did not contribute directly to grain Cd concentrations, it significantly affected PC2.

Overall, the GLS models explained 75–91% of the variance in grain elemental concentrations, except for Fe where the explanatory power was lower (64–70%; Table S5).

To delve further into the effects of climate condition on elemental translocation to grains, retention fractions of Fe, Zn, and Cd in straw and husks were calculated (i.e., from the total elemental contents in each organ; Fig. 4, Table S3). Straw Fe retention averaged approximately 55% and husk Fe retention averaged approximately 84% across soil As and climate condition. Straw Zn retention averaged 30% and husk Zn retention 22%. For both straw and husks, no consistent significant changes in Fe or Zn retention were observed in any treatment compared to low-As and today’s conditions.

**Fig. 4:**
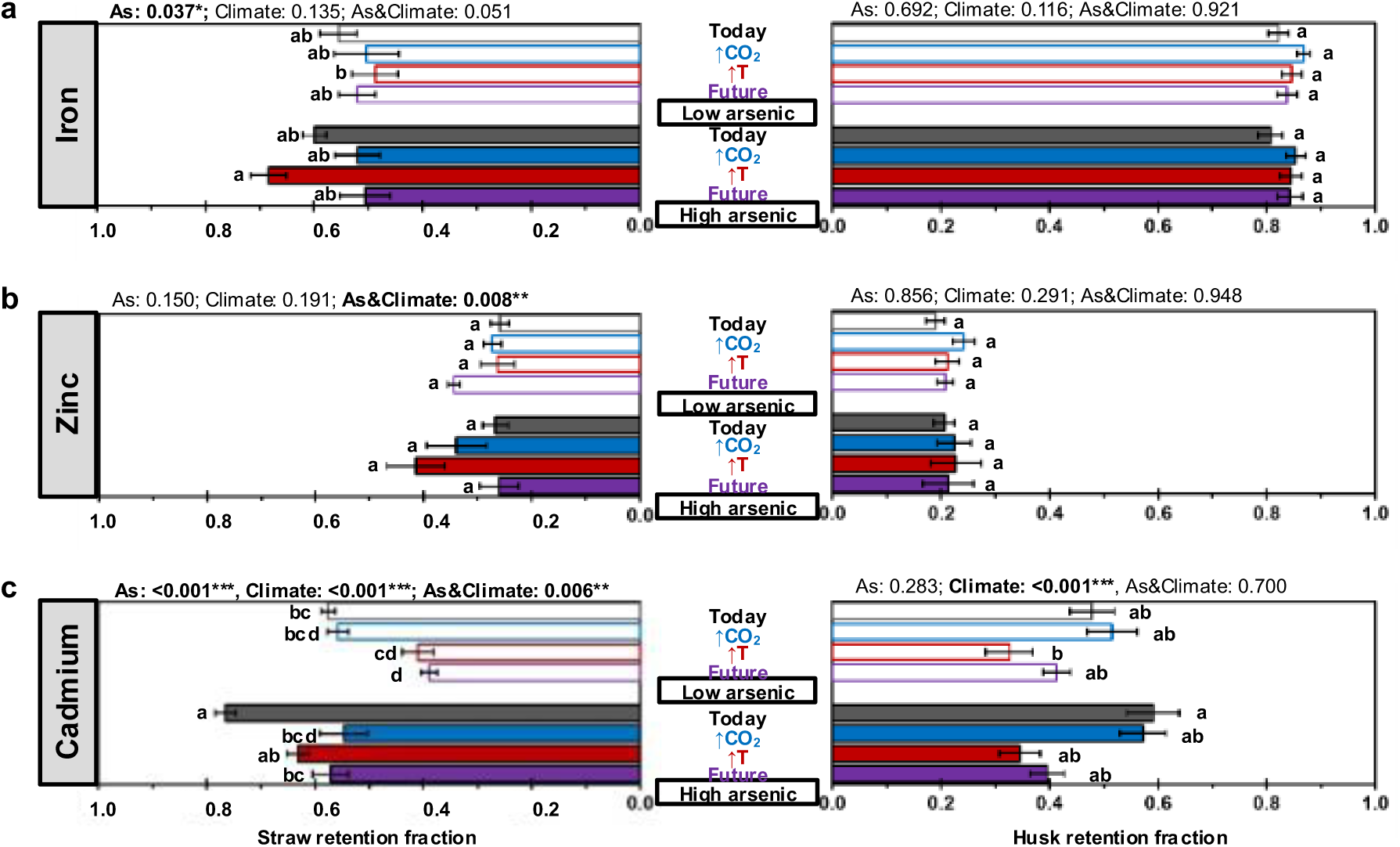
Retention fractions in Oryza sativa L. cv. M206 grown under varying climatic and soil arsenic conditions. Retention fractions indicate the portion of total tissue elemental content that is retained in this tissue relative to the upper tissues (see methods section for precise calculations). Straw (left panels) and husk (right panels) retention fractions for (a) iron, (b) zinc, and (c) cadmium in rice are presented. Rice was subjected to today’s (33°C, 415 ppmv CO2; grey), elevated CO2 (33°C, 850 ppmv CO2; blue), elevated temperature (38°C, 415 ppmv CO2; red), and future (38°C, 850 ppmv CO2; purple) climatic conditions with low (no fill, 7.3 mg As kg^−1^) and high (filled, amended to 24.5 mg As kg^−1^) soil arsenic levels. Bars present the mean ± standard deviation of eight biological replicates for iron and zinc, and four biological replicates for cadmium. Statistical analyses of climate, arsenic, and their interaction on retention fractions were performed using a two-factor ANOVA at a 95% confidence level using the results of a generalized least squares test. Results are shown above each plot, with significant findings highlighted in bold. Post-hoc comparisons using the Emmeans test are indicated by different lowercase letters, where letters denote statistically significant differences (p<0.05). Details of analytical replicates and statistical calculations are provided in Tables S2 and S3, respectively.

Contrary to Fe and Zn, future conditions altered Cd retention in straw in both low-and high-As soils (Fig. 4c). In the low-As soil, straw Cd retention reached 58±4 % under today’s climate, but decreased by 33% to 39±4 % under future conditions. Comparing high-and low-As soils, straw Cd retention increased by 33% to 77±5 % in the high-As soil compared to the low-As soil under today’s climate. Here, future conditions decreased straw Cd retention to levels observed in the low-As soil under today’s climate. An interaction between climate and soil As on straw Cd retention was evident, as elevated CO_2_ in the high-As soil also reduced retention to levels observed in the low-As soil under today’s climate. Husk Cd retention followed the same trends as straw Cd retention, although these effects were less pronounced given lower replicate numbers (grain Cd with only 4 replicates, whereas other tissue had 8 replicates). Soil As did not significantly affect husk Cd retention, whereas climate condition exerted a significant temperature-dependent effect. Husk Cd retention averaged approximately 54% under today’s climate in both soils and decreased by approximately 30% to 32% with increasing temperature.

## Discussion

### Grain micronutrients and Cd under combined arsenic contamination and future climate

The individual and combined effects of soil As contamination and future climatic conditions on micronutrients Fe and Zn and the toxin Cd in rice grains were investigated. The results align with previous work showing decreased rice grain Zn and no change in grain Fe with increasing As contamination (Fig. 1a,b, considering pairwise comparison) (Williams et al. 2009; Vromman et al. 2013). Because As contamination led to decreased aboveground Zn content (Fig. 1e, S2) and had no effect on internal translocation (Fig. 4b), decreased grain Zn concentration was likely due to interrupted metabolic pathways e.g. via oxidative stress (Yu et al. 2012; Koley et al. 2023). In contrast to prominent mono-factorial studies investigating the response of micronutrients to elevated CO_2_ with climate change (Myers et al. 2014; Zhu et al. 2018; Ter Haar et al. 2025), our study showed increased concentrations of Zn (and slightly Fe) in temperature-related future scenarios (Fig. 1a,b). This highlights the critical importance of including multiple climate factors, especially elevated temperature, to investigate the effect of climate change on grain nutritional quality. When considered in combination, As contamination and future climate had only little effect on Fe grain concentrations, which follows as their individual effects were also low (Fig. 1a). The counteracting effects of As contamination (which decreased grain Zn) and future climate (which increased grain Zn) led to a significant 25% decrease in grain Zn concentrations in the high-As soil under future conditions compared to low-As soil under today’s conditions (Fig. 1b). Together, our results indicate that certain micronutrients (e.g., Zn) decrease in rice grains with introduction of As contamination and increase under future climatic conditions (i.e., in the long term), with combined As contamination and climate change, leading to a slight decrease in grain concentrations.

Cadmium uptake and translocation to rice grains followed similar trends as Fe and Zn in its response to soil As contamination, but very different trends in its response to future climate. Similar to Fe and Zn, a tendency for reduced grain Cd was observed with As contamination and supported by a significant As term in the ANOVA (Fig. 1c). A low replication of 4 for Cd grain compared to 8 for Zn and Fe reduced statistical sensitivity, which is likely why pairwise comparison was not significant. Contrary to Fe and Zn, grain Cd concentrations increased substantially under future climatic conditions (>three-fold increase, compared to ∼30-60% increase for Zn and no statistically significant increase for Fe). Our results are consistent with the findings of Ge et al. (2015, 2016, 2019) who showed that Cd concentrations in rice shoots and grains increased with higher temperatures. The substantial increase in grain Cd concentrations with future climatic conditions and slight decrease with soil As contamination led to a two-fold increase in grain Cd concentrations compared to low As and today’s conditions. Though this was not statistically significant (given low replication of the grain Cd data), it is still noteworthy as it is in direct contrast to the either no change (Fe) or slight decreasing (Zn) trends observed for micronutrients.

### Biomass production outweighs internal translocation for micronutrients but not for cadmium

Biomass production, on top of internal translocation, was the dominant control of grain Fe and Zn concentrations. This was illustrated by PCA analyses showing strong positive correlations between grain Fe and Zn (i.e, primarily driven by biomass allocation; Fig. 3a,b). For future climate, decreased biomass production due to the stress of elevated temperature likely concentrated grain Zn (and slightly Fe). Arsenic toxicity interrupted both biomass production and metabolic pathways of Zn uptake. We also observed a positive correlation of grain Fe and Zn with PC2 (i.e., elemental concentrations of straw, panicle, and husks), indicating that internal translocation is also important for grain concentrations of these elements. Yet, we did not observe any significant difference in retention fractions of Fe and Zn in straw or husks with future climate compared to today, neither in low-nor high-As soils (Fig. 4a,b). This further supports the outsized importance of biomass production in controlling grain micronutrient responses to future climate and As contamination.

Grain Cd was controlled more by internal translocation than biomass, in contrast to Fe and Zn. There was consistently lower Cd retention in straw in temperature-related climate conditions compared to today, both in low-and high-As soils (Fig. 4c). Husk retention of Cd also decreased with climate change, as future climate conditions markedly reduced the efficacy of the husk barrier for Cd (Fig. 4c). Future climate may have led to saturation of Cd storage mechanisms within straw (Cao et al. 2018; Wiggenhauser et al. 2021) and husks (Yu et al. 2016), or to bypassing of these storage mechanisms via xylem transport (Uraguchi et al. 2009). In the high-As soil, the role of internal translocation became more pronounced. PCA revealed grain Cd only correlated with PC2, which was driven primarily by straw and husk Cd concentrations and only minor contributions of biomass. There was no significant correlation between grain Cd in the high-As soil and the mainly biomass-driven PC1 (Table S5). Together, our findings suggest that high soil As exacerbates the climate-induced overwhelming of sequestration pathways that mitigate Cd transfer to grains. This result parallels the findings of Muehe et al. (2019), who showed that combined high soil As and future climate saturated husk As retention, leading to higher concentrations of grain As. Cadmium and As are both toxic elements that accumulate in rice grains, and management strategies are often sought to mitigate both elements simultaneously (Zhao and Wang 2020; Yin et al. 2025). Our results and those of Muehe et al. (2019) indicate that strategies fortifying Cd and As sequestration mechanisms in straw and husks, e.g. via Si application (Ma et al. 2015; Yu et al. 2016; Limmer and Seyfferth 2022), may be an essential focus to preserve safe grain production in the face of increasing soil As contamination combined with climate change.

Porewater elemental concentrations did not appreciably influence grain micronutrients or Cd(in contrast to biomass and elemental tissue concentrations). Although soil As and climate conditions affected porewater Fe concentrations (Fig. 2a), this translated only into differences between the elevated CO₂ and elevated temperature treatments in grain Fe concentrations (Fig. 1a) and was not reflected in the PCA-based predictors identified by the statistical models (Table S5). Plants were likely not Fe-limited, as Fe is often abundant in flooded paddy soils due to large concentrations of reduced Fe^2+^ (Tinh et al. 2001). In these systems, Fe uptake is regulated primarily by physiological mechanisms that prevent excessive accumulation and toxicity (Becker and Asch 2005; Kirk et al. 2022). Overall, this contrasts with the findings for As, as porewater As concentrations determined subsequent grain concentrations (Muehe et al. (2019). While not significant controllers of grain Fe, Zn, and Cd, our detailed analysis of porewater chemistry provides key insights into dynamics of solid-liquid interactions, which are detailed in Note S1.

### Societal implications and future work

Our work illustrates the complex interactions that result from the combined effects of As contamination and climate change on metal concentrations in rice grains. We find that while grain Fe is likely to experience no change, grain Zn concentrations decrease by 25% with these combined stressors, contributing to human Zn deficiency (Liu et al. 2024). Even more concerning is the observed increasing trend in rice grain Cd concentrations observed with combined As contamination and future climatic conditions that has wide-reaching deleterious effects on human health (Qu and Zheng 2024). Thus, considering the combined threat of soil As contamination and a changing climate, deteriorating nutritional quality of produced rice grains and threatened human health can be expected and must be mitigated in the coming decades and centuries. Further investigation is urgently needed for a wider variety of rice cultivars to more thoroughly establish the potential implications of internal translocation mechanisms on supply of micronutrient vs. toxic metals from rice-derived food products. Moreover, our results provide evidence that strategies such as plant breeding and soil amendment with inputs should be prioritized in rice cultivation, as these have shown considerable potential to mitigate Cd accumulation in rice while maintaining or enhancing micronutrients such as Fe and Zn (Ueno et al. 2010; Swamy et al. 2016; Cao et al. 2018; Viana et al. 2022; Tan et al. 2023).

## Data availability

The data that support the findings of this study are published in the main text and supplement, with additional data published in Pangaea: https://www.pangaea.de/?t=Agriculture

## Supporting information

*File 1:* Supplementary data, statistical outputs, additional discussion.

## Acknowledgements

The authors are deeply grateful to Carnegie greenhouse staff for greenhouse work and support. Additionally, acknowledgements go to Doug Turner and Guangchao Li for technical lab support, to the Soil and Environmental Biogeochemistry research group for data discussion and to interns in the lab in 2016 and 2017. This work was funded by Marie Skłodowska-Curie Postdoctoral Fellowship (RACe), the Helmholtz Young Investigator Grant RhizoThreats, and by the Deutsche Forschungsgemeinschaft (DFG, German Research Foundation) under Germany’s Excellence Strategy -EXC 3121 – 533771652.

## Statements and declarations

### Funding

This work was funded by Marie Skłodowska-Curie Postdoctoral Fellowship (RACe), the Helmholtz Young Investigator Grant RhizoThreats, and by the Deutsche Forschungsgemeinschaft (DFG, German Research Foundation) under Germany’s Excellence Strategy -EXC 3121 – 533771652.

### Competing interests

The authors have no relevant financial or non-financial interests to disclose.

### Author contributions

Funding was acquired by E. Marie Muehe. The study was conceptualized by E. Marie Muehe and Scott Fendorf. The experiment, data acquisition, and data analyses were performed by E. Marie Muehe and Tianmei Wang. Statistical analyses were performed by E. Marie Muehe and Jill Bachelder. The data was conceptualized for publishing and then written by Jill Bachelder and E. Marie Muehe, with input from Scott Fendorf and Tianmei Wang. All authors have read and agreed to the final manuscript.

## Notes

### Competing Interest Statement

The authors have declared no competing interest.

