## Supplementary file for "Combined climate change and arsenic pollution increases rice cadmium with little effect on micronutrients"

*Corresponding author

**Table S1** Elemental contents of Californian paddy soil.

| Element | Concen | tration | Unit |
| --- | --- | --- | --- |
| P^‖^ | 428.00 | ±4.29 | mg kg^-1^ dry soil |
| K^‖^ | 10.95 | ±0.01 | g kg^-1^ dry soil |
| Fe^‖^ | 54.44 | ±0.02 | g kg^-1^ dry soil |
| S^‖^ | 373.69 | ±2.50 | mg kg^-1^ dry soil |
| Zn^‖^ | 126.98 | ±0.35 | mg kg^-1^ dry soil |
| Cd^‖^ | 2.91 | ±0.11 | mg kg^-1^ dry soil |
| Mn^‖^ | 789.88 | ±0.53 | mg kg^-1^ dry soil |
| Ni^‖^ | 136.43 | ±0.51 | mg kg^-1^ dry soil |
| Cr^‖^ | 173.74 | ±0.32 | mg kg^-1^ dry soil |
| Cu^‖^ | 63.79 | ±0.35 | mg kg^-1^ dry soil |
| ^‖^ Determined by X-ray fluorescence (Spectro XePositive HE XRF Spectrometer, AMETEK, Kleve, Germany) | | | |

**Table S2** List of excluded datapoints in grain, husk, panicle, and straw datasets.

| Tissue | As, Climate, Replicate | Element | Reason |
| --- | --- | --- | --- |
| Grain | All, every second sample | Cd | Below quantification limit of ICP-MS |
| Husk | Low As, eCO_2_, 4 | All elements | Sample spilled |
|  | Low As, Future, 2  High As, Today, 8 | Cd  Cd | Cd was not properly measured in the ICP-MS |
| Panicle | Low As, Today, 5 | All elements | Sample spilled |
|  | Low As, Today, 6 | Cd, Zn, Cu, Mn, Ni, Cr | Sample for ICP-MS not aligned |
|  | Low As, eCO_2_, 2  Low As, eT, 8  Low As, Future, 2  High As, Today, 2  High As, eCO_2_, 2  High As, eCO_2_, 6  High As, eT, 4  High As, eT, 6  High As, Future, 6 | Cr  Cr  Cr  Cr  Cr  Cr  Cr  Cr  Cr | Below quantification limit of ICP-MS |
|  | High As, Today, 7 | Fe, K, P, S | Sample spilled for ICP-OES |
|  | High As, Future, 6  High As, Future, 7 | K  K | K was not properly measured in ICP-OES |
| Straw | All treatments | K | K above the calibration of the ICP-OES |

**Table S3** Summaries of generalized linear models.

Values provided are for climate conditions of today (33ºC, 415 ppmv CO_2_), elevated CO_2_ (eCO_2_; 33ºC, 850 ppmv CO_2_), elevated temperature (eT; 38ºC, 415 ppmv CO_2_) and future (38ºC, 850 ppmv CO_2_) paired with low (no fill, 7.3 mg As kg^−1^dry soil) and high (filled, amended to 24.5 mg As kg^−1^dry soil) soil arsenic levels. Post-hoc comparisons were performed using the Emmeans test. For samples whose residuals were shown by Levene and Shapiro-Wilkes tests to be of non-homogenous variance, a constant variance function was employed to eliminate the effect of heteroscedasticity from the low- vs. high-As treatments (“varIdent” indicated in subsection headings, where applicable).

|  | **Estimate** | **Std. Error** | **df** | **t value** | **Pr(>\|t\|)** |
| --- | --- | --- | --- | --- | --- |
| **SECTION A – GRAIN BIOMASS** | | |  |  |  |
| **Grain weight per plant** |  |  |  |  |  |
| Low Today - Low Future | 1.82575 | 1.058585 | 56 | 1.724708 | 0.671485 |
| Low Today - Low eT | 1.505875 | 1.058585 | 56 | 1.422535 | 0.842984 |
| Low Today - Low eCO2 | -0.97088 | 1.058585 | 56 | -0.91714 | 0.983139 |
| Low Today - High Today | 4.52525 | 1.058585 | 56 | 4.274809 | 0.001814 |
| Low Today - High Future | 4.822125 | 1.058585 | 56 | 4.555254 | 0.000716 |
| Low Today - High eT | 7.500375 | 1.058585 | 56 | 7.085282 | 6.87E-08 |
| Low Today - High eCO2 | 2.74375 | 1.058585 | 56 | 2.591903 | 0.180681 |
| Low Future - High Future | 2.996375 | 1.058585 | 56 | 2.830547 | 0.107958 |
| Low eT - Low Future | 0.319875 | 1.058585 | 56 | 0.302172 | 0.999987 |
| Low eT - Low eCO2 | -2.47675 | 1.058585 | 56 | -2.33968 | 0.291302 |
| Low eT - High Future | 3.31625 | 1.058585 | 56 | 3.132719 | 0.052027 |
| Low eT - High eT | 5.9945 | 1.058585 | 56 | 5.662747 | 1.42E-05 |
| Low eT - High eCO2 | 1.237875 | 1.058585 | 56 | 1.169367 | 0.937243 |
| Low eCO2 - Low Future | 2.796625 | 1.058585 | 56 | 2.641851 | 0.163007 |
| Low eCO2 - High Future | 5.793 | 1.058585 | 56 | 5.472398 | 2.85E-05 |
| Low eCO2 - High eCO2 | 3.714625 | 1.058585 | 56 | 3.509047 | 0.018879 |
| High Today - Low Future | -2.6995 | 1.058585 | 56 | -2.5501 | 0.196531 |
| High Today - Low eT | -3.01937 | 1.058585 | 56 | -2.85227 | 0.102723 |
| High Today - Low eCO2 | -5.49612 | 1.058585 | 56 | -5.19195 | 7.82E-05 |
| High Today - High Future | 0.296875 | 1.058585 | 56 | 0.280445 | 0.999992 |
| High Today - High eT | 2.975125 | 1.058585 | 56 | 2.810473 | 0.112986 |
| High Today - High eCO2 | -1.7815 | 1.058585 | 56 | -1.68291 | 0.69791 |
| High eT - Low Future | -5.67462 | 1.058585 | 56 | -5.36057 | 4.27E-05 |
| High eT - Low eCO2 | -8.47125 | 1.058585 | 56 | -8.00243 | 2.13E-09 |
| High eT - High Future | -2.67825 | 1.058585 | 56 | -2.53003 | 0.204491 |
| High eT - High eCO2 | -4.75663 | 1.058585 | 56 | -4.49338 | 0.000881 |
| High eCO2 - Low Future | -0.918 | 1.058585 | 56 | -0.8672 | 0.987812 |
| High eCO2 - High Future | 2.078375 | 1.058585 | 56 | 1.963352 | 0.515001 |
| **SECTION B – GRAIN ELEMENTAL CONCENTRATIONS** | | | |  |  |
| **Grain Fe concentration** |  |  |  |  |  |
| Low Today - Low Future | -1.25625 | 0.78457 | 56 | -1.6012 | 0.74756 |
| Low Today - Low eT | -1.94 | 0.78457 | 56 | -2.47269 | 0.228483 |
| Low Today - Low eCO2 | 1.5025 | 0.78457 | 56 | 1.915062 | 0.546766 |
| Low Today - High Today | 1.99875 | 0.78457 | 56 | 2.547575 | 0.19752 |
| Low Today - High Future | 0.78 | 0.78457 | 56 | 0.994175 | 0.97344 |
| Low Today - High eT | -0.85875 | 0.78457 | 56 | -1.09455 | 0.955361 |
| Low Today - High eCO2 | 4.045 | 0.78457 | 56 | 5.155692 | 8.90E-05 |
| Low Future - High Future | 2.03625 | 0.78457 | 56 | 2.595371 | 0.179409 |
| Low eT - Low Future | 0.68375 | 0.78457 | 56 | 0.871497 | 0.987454 |
| Low eT - Low eCO2 | 3.4425 | 0.78457 | 56 | 4.387755 | 0.001253 |
| Low eT - High Future | 2.72 | 0.78457 | 56 | 3.466868 | 0.02126 |
| Low eT - High eT | 1.08125 | 0.78457 | 56 | 1.378144 | 0.863219 |
| Low eT - High eCO2 | 5.985 | 0.78457 | 56 | 7.628385 | 8.75E-09 |
| Low eCO2 - Low Future | -2.75875 | 0.78457 | 56 | -3.51626 | 0.018497 |
| Low eCO2 - High Future | -0.7225 | 0.78457 | 56 | -0.92089 | 0.982741 |
| Low eCO2 - High eCO2 | 2.5425 | 0.78457 | 56 | 3.24063 | 0.039336 |
| High Today - Low Future | -3.255 | 0.78457 | 56 | -4.14877 | 0.002725 |
| High Today - Low eT | -3.93875 | 0.78457 | 56 | -5.02027 | 0.000144 |
| High Today - Low eCO2 | -0.49625 | 0.78457 | 56 | -0.63251 | 0.998242 |
| High Today - High Future | -1.21875 | 0.78457 | 56 | -1.5534 | 0.775063 |
| High Today - High eT | -2.8575 | 0.78457 | 56 | -3.64212 | 0.012877 |
| High Today - High eCO2 | 2.04625 | 0.78457 | 56 | 2.608117 | 0.174794 |
| High eT - Low Future | -0.3975 | 0.78457 | 56 | -0.50665 | 0.999583 |
| High eT - Low eCO2 | 2.36125 | 0.78457 | 56 | 3.009611 | 0.070738 |
| High eT - High Future | 1.63875 | 0.78457 | 56 | 2.088724 | 0.434592 |
| High eT - High eCO2 | 4.90375 | 0.78457 | 56 | 6.250241 | 1.61E-06 |
| High eCO2 - Low Future | -5.30125 | 0.78457 | 56 | -6.75689 | 2.38E-07 |
| High eCO2 - High Future | -3.265 | 0.78457 | 56 | -4.16152 | 0.002616 |
| **Grain Zn concentration** |  |  |  |  |  |
| Low Today - Low Future | -9.45625 | 1.640135 | 56 | -5.76553 | 9.75E-06 |
| Low Today - Low eT | -8.59375 | 1.640135 | 56 | -5.23966 | 6.60E-05 |
| Low Today - Low eCO2 | -0.4575 | 1.640135 | 56 | -0.27894 | 0.999993 |
| Low Today - High Today | 15.4 | 1.640135 | 56 | 9.389473 | 1.92E-11 |
| Low Today - High Future | 7.455 | 1.640135 | 56 | 4.545359 | 0.00074 |
| Low Today - High eT | 6.91625 | 1.640135 | 56 | 4.216879 | 0.00219 |
| Low Today - High eCO2 | 17.82 | 1.640135 | 56 | 10.86496 | 7.41E-12 |
| Low Future - High Future | 16.91125 | 1.640135 | 56 | 10.31089 | 7.78E-12 |
| Low eT - Low Future | -0.8625 | 1.640135 | 56 | -0.52587 | 0.999468 |
| Low eT - Low eCO2 | 8.13625 | 1.640135 | 56 | 4.960721 | 0.000177 |
| Low eT - High Future | 16.04875 | 1.640135 | 56 | 9.78502 | 1.02E-11 |
| Low eT - High eT | 15.51 | 1.640135 | 56 | 9.456541 | 1.66E-11 |
| Low eT - High eCO2 | 26.41375 | 1.640135 | 56 | 16.10462 | 7.31E-12 |
| Low eCO2 - Low Future | -8.99875 | 1.640135 | 56 | -5.48659 | 2.71E-05 |
| Low eCO2 - High Future | 7.9125 | 1.640135 | 56 | 4.824299 | 0.000285 |
| Low eCO2 - High eCO2 | 18.2775 | 1.640135 | 56 | 11.1439 | 7.36E-12 |
| High Today - Low Future | -24.8563 | 1.640135 | 56 | -15.155 | 7.31E-12 |
| High Today - Low eT | -23.9938 | 1.640135 | 56 | -14.6291 | 7.31E-12 |
| High Today - Low eCO2 | -15.8575 | 1.640135 | 56 | -9.66841 | 1.16E-11 |
| High Today - High Future | -7.945 | 1.640135 | 56 | -4.84411 | 0.000266 |
| High Today - High eT | -8.48375 | 1.640135 | 56 | -5.17259 | 8.38E-05 |
| High Today - High eCO2 | 2.42 | 1.640135 | 56 | 1.475489 | 0.816917 |
| High eT - Low Future | -16.3725 | 1.640135 | 56 | -9.98241 | 8.73E-12 |
| High eT - Low eCO2 | -7.37375 | 1.640135 | 56 | -4.49582 | 0.000874 |
| High eT - High Future | 0.53875 | 1.640135 | 56 | 0.328479 | 0.999977 |
| High eT - High eCO2 | 10.90375 | 1.640135 | 56 | 6.648082 | 3.60E-07 |
| High eCO2 - Low Future | -27.2763 | 1.640135 | 56 | -16.6305 | 7.31E-12 |
| High eCO2 - High Future | -10.365 | 1.640135 | 56 | -6.3196 | 1.24E-06 |
| **Grain Cd concentration** |  |  |  |  |  |
| Low Today - Low Future | -7.9975 | 1.401585 | 24 | -5.70604 | 0.000166 |
| Low Today - Low eT | -8.5525 | 1.401585 | 24 | -6.10202 | 6.36E-05 |
| Low Today - Low eCO2 | 0.34 | 1.401585 | 24 | 0.242583 | 0.999997 |
| Low Today - High Today | 2.235 | 1.401585 | 24 | 1.594623 | 0.748847 |
| Low Today - High Future | -4.1225 | 1.401585 | 24 | -2.94131 | 0.107791 |
| Low Today - High eT | -6.21 | 1.401585 | 24 | -4.4307 | 0.00374 |
| Low Today - High eCO2 | -1.23 | 1.401585 | 24 | -0.87758 | 0.985439 |
| Low Future - High Future | 3.875 | 1.401585 | 24 | 2.764727 | 0.151439 |
| Low eT - Low Future | 0.555 | 1.401585 | 24 | 0.39598 | 0.999905 |
| Low eT - Low eCO2 | 8.8925 | 1.401585 | 24 | 6.344603 | 3.56E-05 |
| Low eT - High Future | 4.43 | 1.401585 | 24 | 3.160708 | 0.068951 |
| Low eT - High eT | 2.3425 | 1.401585 | 24 | 1.671322 | 0.704381 |
| Low eT - High eCO2 | 7.3225 | 1.401585 | 24 | 5.224443 | 0.000538 |
| Low eCO2 - Low Future | -8.3375 | 1.401585 | 24 | -5.94862 | 9.21E-05 |
| Low eCO2 - High Future | -4.4625 | 1.401585 | 24 | -3.1839 | 0.065678 |
| Low eCO2 - High eCO2 | -1.57 | 1.401585 | 24 | -1.12016 | 0.945859 |
| High Today - Low Future | -10.2325 | 1.401585 | 24 | -7.30066 | 3.82E-06 |
| High Today - Low eT | -10.7875 | 1.401585 | 24 | -7.69664 | 1.56E-06 |
| High Today - Low eCO2 | -1.895 | 1.401585 | 24 | -1.35204 | 0.869345 |
| High Today - High Future | -6.3575 | 1.401585 | 24 | -4.53594 | 0.002897 |
| High Today - High eT | -8.445 | 1.401585 | 24 | -6.02532 | 7.66E-05 |
| High Today - High eCO2 | -3.465 | 1.401585 | 24 | -2.4722 | 0.253933 |
| High eT - Low Future | -1.7875 | 1.401585 | 24 | -1.27534 | 0.899174 |
| High eT - Low eCO2 | 6.55 | 1.401585 | 24 | 4.673281 | 0.002074 |
| High eT - High Future | 2.0875 | 1.401585 | 24 | 1.489385 | 0.805486 |
| High eT - High eCO2 | 4.98 | 1.401585 | 24 | 3.553121 | 0.029387 |
| High eCO2 - Low Future | -6.7675 | 1.401585 | 24 | -4.82846 | 0.001419 |
| High eCO2 - High Future | -2.8925 | 1.401585 | 24 | -2.06374 | 0.463819 |
| **SECTION C – GRAIN ELEMENTAL CONTENTS** | |  |  |  |  |
| **Grain Fe content** |  |  |  |  |  |
| Low Today - Low Future | 21.50125 | 17.63151 | 56 | 1.219479 | 0.922688 |
| Low Today - Low eT | 8.4575 | 17.63151 | 56 | 0.479681 | 0.99971 |
| Low Today - Low eCO2 | 2.1725 | 17.63151 | 56 | 0.123217 | 1 |
| Low Today - High Today | 97.53125 | 17.63151 | 56 | 5.531644 | 2.30E-05 |
| Low Today - High Future | 93.545 | 17.63151 | 56 | 5.305557 | 5.21E-05 |
| Low Today - High eT | 137.075 | 17.63151 | 56 | 7.774433 | 5.03E-09 |
| Low Today - High eCO2 | 84.735 | 17.63151 | 56 | 4.805884 | 0.000304 |
| Low Future - High Future | 72.04375 | 17.63151 | 56 | 4.086079 | 0.003327 |
| Low eT - Low Future | 13.04375 | 17.63151 | 56 | 0.739798 | 0.99531 |
| Low eT - Low eCO2 | -6.285 | 17.63151 | 56 | -0.35646 | 0.999961 |
| Low eT - High Future | 85.0875 | 17.63151 | 56 | 4.825877 | 0.000284 |
| Low eT - High eT | 128.6175 | 17.63151 | 56 | 7.294752 | 3.10E-08 |
| Low eT - High eCO2 | 76.2775 | 17.63151 | 56 | 4.326203 | 0.001534 |
| Low eCO2 - Low Future | 19.32875 | 17.63151 | 56 | 1.096262 | 0.954992 |
| Low eCO2 - High Future | 91.3725 | 17.63151 | 56 | 5.182341 | 8.10E-05 |
| Low eCO2 - High eCO2 | 82.5625 | 17.63151 | 56 | 4.682667 | 0.000465 |
| High Today - Low Future | -76.03 | 17.63151 | 56 | -4.31217 | 0.001606 |
| High Today - Low eT | -89.0737 | 17.63151 | 56 | -5.05196 | 0.000129 |
| High Today - Low eCO2 | -95.3587 | 17.63151 | 56 | -5.40843 | 3.59E-05 |
| High Today - High Future | -3.98625 | 17.63151 | 56 | -0.22609 | 0.999998 |
| High Today - High eT | 39.54375 | 17.63151 | 56 | 2.242788 | 0.343119 |
| High Today - High eCO2 | -12.7963 | 17.63151 | 56 | -0.72576 | 0.995832 |
| High eT - Low Future | -115.574 | 17.63151 | 56 | -6.55495 | 5.11E-07 |
| High eT - Low eCO2 | -134.903 | 17.63151 | 56 | -7.65122 | 8.03E-09 |
| High eT - High Future | -43.53 | 17.63151 | 56 | -2.46888 | 0.230146 |
| High eT - High eCO2 | -52.34 | 17.63151 | 56 | -2.96855 | 0.078141 |
| High eCO2 - Low Future | -63.2337 | 17.63151 | 56 | -3.58641 | 0.015136 |
| High eCO2 - High Future | 8.81 | 17.63151 | 56 | 0.499674 | 0.99962 |
| **Grain Zn content** |  |  |  |  |  |
| Low Today - Low Future | -37.5213 | 27.21412 | 56 | -1.37874 | 0.862957 |
| Low Today - Low eT | -40.9825 | 27.21412 | 56 | -1.50593 | 0.801036 |
| Low Today - Low eCO2 | -32.9338 | 27.21412 | 56 | -1.21017 | 0.925543 |
| Low Today - High Today | 233.985 | 27.21412 | 56 | 8.597925 | 2.32E-10 |
| Low Today - High Future | 186.2525 | 27.21412 | 56 | 6.843965 | 1.71E-07 |
| Low Today - High eT | 244.2925 | 27.21412 | 56 | 8.976681 | 6.20E-11 |
| Low Today - High eCO2 | 229.7125 | 27.21412 | 56 | 8.440929 | 4.13E-10 |
| Low Future - High Future | 223.7738 | 27.21412 | 56 | 8.222706 | 9.30E-10 |
| Low eT - Low Future | 3.46125 | 27.21412 | 56 | 0.127186 | 1 |
| Low eT - Low eCO2 | 8.04875 | 27.21412 | 56 | 0.295756 | 0.999989 |
| Low eT - High Future | 227.235 | 27.21412 | 56 | 8.349892 | 5.79E-10 |
| Low eT - High eT | 285.275 | 27.21412 | 56 | 10.48261 | 7.59E-12 |
| Low eT - High eCO2 | 270.695 | 27.21412 | 56 | 9.946857 | 8.92E-12 |
| Low eCO2 - Low Future | -4.5875 | 27.21412 | 56 | -0.16857 | 1 |
| Low eCO2 - High Future | 219.1863 | 27.21412 | 56 | 8.054136 | 1.75E-09 |
| Low eCO2 - High eCO2 | 262.6463 | 27.21412 | 56 | 9.651101 | 1.19E-11 |
| High Today - Low Future | -271.506 | 27.21412 | 56 | -9.97667 | 8.76E-12 |
| High Today - Low eT | -274.968 | 27.21412 | 56 | -10.1039 | 8.25E-12 |
| High Today - Low eCO2 | -266.919 | 27.21412 | 56 | -9.8081 | 9.93E-12 |
| High Today - High Future | -47.7325 | 27.21412 | 56 | -1.75396 | 0.652679 |
| High Today - High eT | 10.3075 | 27.21412 | 56 | 0.378756 | 0.99994 |
| High Today - High eCO2 | -4.2725 | 27.21412 | 56 | -0.157 | 1 |
| High eT - Low Future | -281.814 | 27.21412 | 56 | -10.3554 | 7.72E-12 |
| High eT - Low eCO2 | -277.226 | 27.21412 | 56 | -10.1869 | 8.02E-12 |
| High eT - High Future | -58.04 | 27.21412 | 56 | -2.13272 | 0.407482 |
| High eT - High eCO2 | -14.58 | 27.21412 | 56 | -0.53575 | 0.999399 |
| High eCO2 - Low Future | -267.234 | 27.21412 | 56 | -9.81967 | 9.83E-12 |
| High eCO2 - High Future | -43.46 | 27.21412 | 56 | -1.59696 | 0.750045 |
| **Grain Cd content** |  |  |  |  |  |
| Low Today - Low Future | -61.37 | 14.03984 | 24 | -4.37113 | 0.004319 |
| Low Today - Low eT | -66.645 | 14.03984 | 24 | -4.74685 | 0.001733 |
| Low Today - Low eCO2 | 2.0825 | 14.03984 | 24 | 0.148328 | 1 |
| Low Today - High Today | 34.1425 | 14.03984 | 24 | 2.431829 | 0.27131 |
| Low Today - High Future | -5.525 | 14.03984 | 24 | -0.39352 | 0.999909 |
| Low Today - High eT | 3.76 | 14.03984 | 24 | 0.267809 | 0.999993 |
| Low Today - High eCO2 | -3.595 | 14.03984 | 24 | -0.25606 | 0.999995 |
| Low Future - High Future | 55.845 | 14.03984 | 24 | 3.977608 | 0.01105 |
| Low eT - Low Future | 5.275 | 14.03984 | 24 | 0.375716 | 0.999933 |
| Low eT - Low eCO2 | 68.7275 | 14.03984 | 24 | 4.895175 | 0.001206 |
| Low eT - High Future | 61.12 | 14.03984 | 24 | 4.353324 | 0.004509 |
| Low eT - High eT | 70.405 | 14.03984 | 24 | 5.014657 | 0.0009 |
| Low eT - High eCO2 | 63.05 | 14.03984 | 24 | 4.49079 | 0.003233 |
| Low eCO2 - Low Future | -63.4525 | 14.03984 | 24 | -4.51946 | 0.003016 |
| Low eCO2 - High Future | -7.6075 | 14.03984 | 24 | -0.54185 | 0.999245 |
| Low eCO2 - High eCO2 | -5.6775 | 14.03984 | 24 | -0.40438 | 0.99989 |
| High Today - Low Future | -95.5125 | 14.03984 | 24 | -6.80296 | 1.21E-05 |
| High Today - Low eT | -100.788 | 14.03984 | 24 | -7.17868 | 5.05E-06 |
| High Today - Low eCO2 | -32.06 | 14.03984 | 24 | -2.2835 | 0.341893 |
| High Today - High Future | -39.6675 | 14.03984 | 24 | -2.82535 | 0.135047 |
| High Today - High eT | -30.3825 | 14.03984 | 24 | -2.16402 | 0.405929 |
| High Today - High eCO2 | -37.7375 | 14.03984 | 24 | -2.68789 | 0.174499 |
| High eT - Low Future | -65.13 | 14.03984 | 24 | -4.63894 | 0.002255 |
| High eT - Low eCO2 | -1.6775 | 14.03984 | 24 | -0.11948 | 1 |
| High eT - High Future | -9.285 | 14.03984 | 24 | -0.66133 | 0.997322 |
| High eT - High eCO2 | -7.355 | 14.03984 | 24 | -0.52387 | 0.999394 |
| High eCO2 - Low Future | -57.775 | 14.03984 | 24 | -4.11507 | 0.00798 |
| High eCO2 - High Future | -1.93 | 14.03984 | 24 | -0.13747 | 1 |
| Low Today - Low Future | -61.37 | 14.03984 | 24 | -4.37113 | 0.004319 |
| **SECTION D – HUSK, PANICLE, AND STRAW ELEMENTAL CONCENTRATIONS** | | | |  |  |
| **Husk Fe concentration** |  |  |  |  |  |
| Low Today - Low Future | -0.01625 | 0.019085 | 55 | -0.85144 | 0.989038 |
| Low Today - Low eT | -0.03 | 0.019085 | 55 | -1.5719 | 0.764524 |
| Low Today - Low eCO2 | -0.01679 | 0.019755 | 55 | -0.84969 | 0.989171 |
| Low Today - High Today | 0.0125 | 0.019085 | 55 | 0.654958 | 0.997803 |
| Low Today - High Future | -0.0175 | 0.019085 | 55 | -0.91694 | 0.983135 |
| Low Today - High eT | -0.00875 | 0.019085 | 55 | -0.45847 | 0.999785 |
| Low Today - High eCO2 | -0.00125 | 0.019085 | 55 | -0.0655 | 1 |
| Low Future - High Future | -0.00125 | 0.019085 | 55 | -0.0655 | 1 |
| Low eT - Low Future | 0.01375 | 0.019085 | 55 | 0.720453 | 0.99601 |
| Low eT - Low eCO2 | 0.013214 | 0.019755 | 55 | 0.668906 | 0.99749 |
| Low eT - High Future | 0.0125 | 0.019085 | 55 | 0.654958 | 0.997803 |
| Low eT - High eT | 0.02125 | 0.019085 | 55 | 1.113428 | 0.951132 |
| Low eT - High eCO2 | 0.02875 | 0.019085 | 55 | 1.506402 | 0.800723 |
| Low eCO2 - Low Future | 0.000536 | 0.019755 | 55 | 0.027118 | 1 |
| Low eCO2 - High Future | -0.00071 | 0.019755 | 55 | -0.03616 | 1 |
| Low eCO2 - High eCO2 | 0.015536 | 0.019755 | 55 | 0.786417 | 0.993179 |
| High Today - Low Future | -0.02875 | 0.019085 | 55 | -1.5064 | 0.800723 |
| High Today - Low eT | -0.0425 | 0.019085 | 55 | -2.22686 | 0.35238 |
| High Today - Low eCO2 | -0.02929 | 0.019755 | 55 | -1.48244 | 0.813281 |
| High Today - High Future | -0.03 | 0.019085 | 55 | -1.5719 | 0.764524 |
| High Today - High eT | -0.02125 | 0.019085 | 55 | -1.11343 | 0.951132 |
| High Today - High eCO2 | -0.01375 | 0.019085 | 55 | -0.72045 | 0.99601 |
| High eT - Low Future | -0.0075 | 0.019085 | 55 | -0.39297 | 0.999923 |
| High eT - Low eCO2 | -0.00804 | 0.019755 | 55 | -0.40677 | 0.999903 |
| High eT - High Future | -0.00875 | 0.019085 | 55 | -0.45847 | 0.999785 |
| High eT - High eCO2 | 0.0075 | 0.019085 | 55 | 0.392975 | 0.999923 |
| High eCO2 - Low Future | -0.015 | 0.019085 | 55 | -0.78595 | 0.993203 |
| High eCO2 - High Future | -0.01625 | 0.019085 | 55 | -0.85144 | 0.989038 |
| **Husk Zn concentration (varIdent)** |  |  |  |  |  |
| Low Today - Low Future | -2.75375 | 1.906233 | 55 | -1.4446 | 0.8323 |
| Low Today - Low eT | -3.30375 | 1.906233 | 55 | -1.73313 | 0.666107 |
| Low Today - Low eCO2 | -2.05839 | 1.973139 | 55 | -1.04321 | 0.965401 |
| Low Today - High Today | 2.8825 | 1.906233 | 55 | 1.512144 | 0.797657 |
| Low Today - High Future | -0.88 | 1.906233 | 55 | -0.46164 | 0.999775 |
| Low Today - High eT | 1.49 | 1.906233 | 55 | 0.781646 | 0.993425 |
| Low Today - High eCO2 | 3.3775 | 1.906233 | 55 | 1.771819 | 0.641128 |
| Low Future - High Future | 1.87375 | 1.906233 | 55 | 0.982959 | 0.975026 |
| Low eT - Low Future | 0.55 | 1.906233 | 55 | 0.288527 | 0.999991 |
| Low eT - Low eCO2 | 1.245357 | 1.973139 | 55 | 0.631155 | 0.998263 |
| Low eT - High Future | 2.42375 | 1.906233 | 55 | 1.271487 | 0.905377 |
| Low eT - High eT | 4.79375 | 1.906233 | 55 | 2.514776 | 0.211065 |
| Low eT - High eCO2 | 6.68125 | 1.906233 | 55 | 3.504949 | 0.019268 |
| Low eCO2 - Low Future | -0.69536 | 1.973139 | 55 | -0.35241 | 0.999963 |
| Low eCO2 - High Future | 1.178393 | 1.973139 | 55 | 0.597217 | 0.998781 |
| Low eCO2 - High eCO2 | 5.435893 | 1.973139 | 55 | 2.754947 | 0.128258 |
| High Today - Low Future | -5.63625 | 1.906233 | 55 | -2.95675 | 0.080721 |
| High Today - Low eT | -6.18625 | 1.906233 | 55 | -3.24527 | 0.039105 |
| High Today - Low eCO2 | -4.94089 | 1.973139 | 55 | -2.50408 | 0.215491 |
| High Today - High Future | -3.7625 | 1.906233 | 55 | -1.97379 | 0.508321 |
| High Today - High eT | -1.3925 | 1.906233 | 55 | -0.7305 | 0.995653 |
| High Today - High eCO2 | 0.495 | 1.906233 | 55 | 0.259674 | 0.999995 |
| High eT - Low Future | -4.24375 | 1.906233 | 55 | -2.22625 | 0.352724 |
| High eT - Low eCO2 | -3.54839 | 1.973139 | 55 | -1.79835 | 0.623816 |
| High eT - High Future | -2.37 | 1.906233 | 55 | -1.24329 | 0.914999 |
| High eT - High eCO2 | 1.8875 | 1.906233 | 55 | 0.990173 | 0.973992 |
| High eCO2 - Low Future | -6.13125 | 1.906233 | 55 | -3.21642 | 0.042173 |
| High eCO2 - High Future | -4.2575 | 1.906233 | 55 | -2.23346 | 0.348646 |
| **Husk Cd concentration** |  |  |  |  |  |
| Low Today - Low Future | -2.15409 | 1.292633 | 53 | -1.66644 | 0.708104 |
| Low Today - Low eT | -2.41975 | 1.248802 | 53 | -1.93766 | 0.532256 |
| Low Today - Low eCO2 | 0.285339 | 1.292633 | 53 | 0.220743 | 0.999999 |
| Low Today - High Today | 1.549196 | 1.292633 | 53 | 1.198481 | 0.928833 |
| Low Today - High Future | -0.04525 | 1.248802 | 53 | -0.03623 | 1 |
| Low Today - High eT | -0.25238 | 1.248802 | 53 | -0.20209 | 0.999999 |
| Low Today - High eCO2 | -1.09188 | 1.248802 | 53 | -0.87434 | 0.98715 |
| Low Future - High Future | 2.108839 | 1.292633 | 53 | 1.631429 | 0.729456 |
| Low eT - Low Future | 0.265661 | 1.292633 | 53 | 0.205519 | 0.999999 |
| Low eT - Low eCO2 | 2.705089 | 1.292633 | 53 | 2.092697 | 0.432782 |
| Low eT - High Future | 2.3745 | 1.248802 | 53 | 1.901422 | 0.556092 |
| Low eT - High eT | 2.167375 | 1.248802 | 53 | 1.735563 | 0.664578 |
| Low eT - High eCO2 | 1.327875 | 1.248802 | 53 | 1.063319 | 0.961571 |
| Low eCO2 - Low Future | -2.43943 | 1.335026 | 53 | -1.82725 | 0.604965 |
| Low eCO2 - High Future | -0.33059 | 1.292633 | 53 | -0.25575 | 0.999996 |
| Low eCO2 - High eCO2 | -1.37721 | 1.292633 | 53 | -1.06543 | 0.961161 |
| High Today - Low Future | -3.70329 | 1.335026 | 53 | -2.77394 | 0.123778 |
| High Today - Low eT | -3.96895 | 1.292633 | 53 | -3.07044 | 0.061817 |
| High Today - Low eCO2 | -1.26386 | 1.335026 | 53 | -0.94669 | 0.979712 |
| High Today - High Future | -1.59445 | 1.292633 | 53 | -1.23349 | 0.918052 |
| High Today - High eT | -1.80157 | 1.292633 | 53 | -1.39372 | 0.856065 |
| High Today - High eCO2 | -2.64107 | 1.292633 | 53 | -2.04317 | 0.463921 |
| High eT - Low Future | -1.90171 | 1.292633 | 53 | -1.47119 | 0.818901 |
| High eT - Low eCO2 | 0.537714 | 1.292633 | 53 | 0.415984 | 0.999887 |
| High eT - High Future | 0.207125 | 1.248802 | 53 | 0.165859 | 1 |
| High eT - High eCO2 | -0.8395 | 1.248802 | 53 | -0.67224 | 0.997399 |
| High eCO2 - Low Future | -1.06221 | 1.292633 | 53 | -0.82174 | 0.991087 |
| High eCO2 - High Future | 1.046625 | 1.248802 | 53 | 0.838103 | 0.98998 |
| **Panicle Fe concentration** |  |  |  |  |  |
| Low Today - Low Future | -2.77571 | 7.67056 | 53 | -0.36187 | 0.999956 |
| Low Today - Low eT | -0.19196 | 7.67056 | 53 | -0.02503 | 1 |
| Low Today - Low eCO2 | -0.31196 | 7.67056 | 53 | -0.04067 | 1 |
| Low Today - High Today | -4.62905 | 8.245605 | 53 | -0.5614 | 0.999179 |
| Low Today - High Future | -2.80696 | 7.67056 | 53 | -0.36594 | 0.999952 |
| Low Today - High eT | 9.716786 | 7.67056 | 53 | 1.266763 | 0.906886 |
| Low Today - High eCO2 | | 2.069286 | 7.67056 | 53 | 0.26977 |
| Low Future - High Future | -0.03125 | 7.410465 | 53 | -0.00422 | 1 |
| Low eT - Low Future | -2.58375 | 7.410465 | 53 | -0.34866 | 0.999966 |
| Low eT - Low eCO2 | -0.12 | 7.410465 | 53 | -0.01619 | 1 |
| Low eT - High Future | -2.615 | 7.410465 | 53 | -0.35288 | 0.999963 |
| Low eT - High eT | 9.90875 | 7.410465 | 53 | 1.337129 | 0.880315 |
| Low eT - High eCO2 | 2.26125 | 7.410465 | 53 | 0.305143 | 0.999986 |
| Low eCO2 - Low Future | -2.46375 | 7.410465 | 53 | -0.33247 | 0.999975 |
| Low eCO2 - High Future | -2.495 | 7.410465 | 53 | -0.33669 | 0.999973 |
| Low eCO2 - High eCO2 | | | 2.38125 | 7.410465 | 53 |
| High Today - Low Future | 1.853333 | 8.004217 | 53 | 0.231545 | 0.999998 |
| High Today - Low eT | 4.437083 | 8.004217 | 53 | 0.554343 | 0.999244 |
| High Today - Low eCO2 | 4.317083 | 8.004217 | 53 | 0.539351 | 0.999368 |
| High Today - High Future | 1.822083 | 8.004217 | 53 | 0.22764 | 0.999998 |
| High Today - High eT | 14.34583 | 8.004217 | 53 | 1.792284 | 0.627873 |
| High Today - High eCO2 | 6.698333 | 8.004217 | 53 | 0.836851 | 0.990069 |
| High eT - Low Future | -12.4925 | 7.410465 | 53 | -1.68579 | 0.696083 |
| High eT - Low eCO2 | -10.0288 | 7.410465 | 53 | -1.35332 | 0.873635 |
| High eT - High Future | -12.5238 | 7.410465 | 53 | -1.69001 | 0.693445 |
| High eT - High eCO2 | -7.6475 | 7.410465 | 53 | -1.03199 | 0.967283 |
| High eCO2 - Low Future | -4.845 | 7.410465 | 53 | -0.65381 | 0.997818 |
| High eCO2 - High Future | -4.87625 | 7.410465 | 53 | -0.65802 | 0.997727 |
| **Panicle Zn concentration** |  |  |  |  |  |
| Low Today - Low Future | -10.2084 | 4.882829 | 54 | -2.09068 | 0.433809 |
| Low Today - Low eT | -4.18429 | 4.882829 | 54 | -0.85694 | 0.988596 |
| Low Today - Low eCO2 | -3.47279 | 4.882829 | 54 | -0.71123 | 0.996309 |
| Low Today - High Today | 3.394958 | 4.882829 | 54 | 0.695285 | 0.996795 |
| Low Today - High Future | -2.04067 | 4.882829 | 54 | -0.41793 | 0.999884 |
| Low Today - High eT | 4.687833 | 4.882829 | 54 | 0.960065 | 0.978077 |
| Low Today - High eCO2 | 3.379958 | 4.882829 | 54 | 0.692213 | 0.996882 |
| Low Future - High Future | 8.16775 | 4.520621 | 54 | 1.806776 | 0.618345 |
| Low eT - Low Future | -6.02413 | 4.520621 | 54 | -1.33259 | 0.88223 |
| Low eT - Low eCO2 | 0.7115 | 4.520621 | 54 | 0.15739 | 1 |
| Low eT - High Future | 2.143625 | 4.520621 | 54 | 0.474188 | 0.99973 |
| Low eT - High eT | 8.872125 | 4.520621 | 54 | 1.96259 | 0.515782 |
| Low eT - High eCO2 | 7.56425 | 4.520621 | 54 | 1.673277 | 0.703885 |
| Low eCO2 - Low Future | -6.73563 | 4.520621 | 54 | -1.48998 | 0.809308 |
| Low eCO2 - High Future | 1.432125 | 4.520621 | 54 | 0.316798 | 0.999982 |
| Low eCO2 - High eCO2 | 6.85275 | 4.520621 | 54 | 1.515887 | 0.795587 |
| High Today - Low Future | -13.6034 | 4.520621 | 54 | -3.00918 | 0.071465 |
| High Today - Low eT | -7.57925 | 4.520621 | 54 | -1.67659 | 0.701824 |
| High Today - Low eCO2 | -6.86775 | 4.520621 | 54 | -1.5192 | 0.793798 |
| High Today - High Future | -5.43563 | 4.520621 | 54 | -1.20241 | 0.927741 |
| High Today - High eT | 1.292875 | 4.520621 | 54 | 0.285995 | 0.999991 |
| High Today - High eCO2 | -0.015 | 4.520621 | 54 | -0.00332 | 1 |
| High eT - Low Future | -14.8963 | 4.520621 | 54 | -3.29518 | 0.034504 |
| High eT - Low eCO2 | -8.16063 | 4.520621 | 54 | -1.8052 | 0.619378 |
| High eT - High Future | -6.7285 | 4.520621 | 54 | -1.4884 | 0.810128 |
| High eT - High eCO2 | | -1.30788 | 4.520621 | 54 | -0.28931 |
| High eCO2 - Low Future | -13.5884 | 4.520621 | 54 | -3.00586 | 0.072042 |
| High eCO2 - High Future | -5.42063 | 4.520621 | 54 | -1.19909 | 0.928723 |
| **Panicle Cd concentration** |  |  |  |  |  |
| Low Today - Low Future | -7.65658 | 6.978543 | 54 | -1.09716 | 0.954697 |
| Low Today - Low eT | | -10.9471 | 6.978543 | 54 | -1.56868 |
| Low Today - Low eCO2 | -0.96208 | 6.978543 | 54 | -0.13786 | 1 |
| Low Today - High Today | -8.95071 | 6.978543 | 54 | -1.2826 | 0.901331 |
| Low Today - High Future | -12.4488 | 6.978543 | 54 | -1.78387 | 0.633317 |
| Low Today - High eT | -11.9573 | 6.978543 | 54 | -1.71344 | 0.678672 |
| Low Today - High eCO2 | -5.99933 | 6.978543 | 54 | -0.85968 | 0.98838 |
| Low Future - High Future | -4.79225 | 6.460875 | 54 | -0.74173 | 0.995216 |
| Low eT - Low Future | 3.2905 | 6.460875 | 54 | 0.509296 | 0.999567 |
| Low eT - Low eCO2 | 9.985 | 6.460875 | 54 | 1.545456 | 0.7794 |
| Low eT - High Future | -1.50175 | 6.460875 | 54 | -0.23244 | 0.999998 |
| Low eT - High eT | -1.01025 | 6.460875 | 54 | -0.15636 | 1 |
| Low eT - High eCO2 | 4.94775 | 6.460875 | 54 | 0.765802 | 0.994184 |
| Low eCO2 - Low Future | -6.6945 | 6.460875 | 54 | -1.03616 | 0.966604 |
| Low eCO2 - High Future | -11.4868 | 6.460875 | 54 | -1.77789 | 0.637211 |
| Low eCO2 - High eCO2 | -5.03725 | 6.460875 | 54 | -0.77965 | 0.993514 |
| High Today - Low Future | | 1.294125 | 6.460875 | 54 | 0.200302 |
| High Today - Low eT | -1.99637 | 6.460875 | 54 | -0.30899 | 0.999985 |
| High Today - Low eCO2 | 7.988625 | 6.460875 | 54 | 1.236462 | 0.917162 |
| High Today - High Future | -3.49813 | 6.460875 | 54 | -0.54143 | 0.999353 |
| High Today - High eT | -3.00662 | 6.460875 | 54 | -0.46536 | 0.999762 |
| High Today - High eCO2 | | 2.951375 | 6.460875 | 54 | 0.456807 |
| High eT - Low Future | 4.30075 | 6.460875 | 54 | 0.665661 | 0.997561 |
| High eT - Low eCO2 | 10.99525 | 6.460875 | 54 | 1.70182 | 0.686024 |
| High eT - High Future | -0.4915 | 6.460875 | 54 | -0.07607 | 1 |
| High eT - High eCO2 | 5.958 | 6.460875 | 54 | 0.922166 | 0.982551 |
| High eCO2 - Low Future | -1.65725 | 6.460875 | 54 | -0.25651 | 0.999996 |
| High eCO2 - High Future | -6.4495 | 6.460875 | 54 | -0.99824 | 0.972761 |
| **Straw Fe concentration** |  |  |  |  |  |
| Low Today - Low Future | 0.03825 | 0.015881 | 56 | 2.408569 | 0.257527 |
| Low Today - Low eT | 0.02175 | 0.015881 | 56 | 1.369578 | 0.866947 |
| Low Today - Low eCO2 | 0.031375 | 0.015881 | 56 | 1.975656 | 0.506956 |
| Low Today - High Today | 0.010625 | 0.015881 | 56 | 0.669047 | 0.997492 |
| Low Today - High Future | 0.039 | 0.015881 | 56 | 2.455796 | 0.23591 |
| Low Today - High eT | 0.0065 | 0.015881 | 56 | 0.409299 | 0.999899 |
| Low Today - High eCO2 | 0.037 | 0.015881 | 56 | 2.329858 | 0.296329 |
| Low Future - High Future | 0.00075 | 0.015881 | 56 | 0.047227 | 1 |
| Low eT - Low Future | 0.0165 | 0.015881 | 56 | 1.038991 | 0.966192 |
| Low eT - Low eCO2 | 0.009625 | 0.015881 | 56 | 0.606078 | 0.998662 |
| Low eT - High Future | 0.01725 | 0.015881 | 56 | 1.086217 | 0.957122 |
| Low eT - High eT | -0.01525 | 0.015881 | 56 | -0.96028 | 0.978112 |
| Low eT - High eCO2 | 0.01525 | 0.015881 | 56 | 0.960279 | 0.978112 |
| Low eCO2 - Low Future | 0.006875 | 0.015881 | 56 | 0.432913 | 0.999854 |
| Low eCO2 - High Future | 0.007625 | 0.015881 | 56 | 0.48014 | 0.999708 |
| Low eCO2 - High eCO2 | 0.005625 | 0.015881 | 56 | 0.354201 | 0.999962 |
| High Today - Low Future | 0.027625 | 0.015881 | 56 | 1.739522 | 0.66199 |
| High Today - Low eT | 0.011125 | 0.015881 | 56 | 0.700532 | 0.996654 |
| High Today - Low eCO2 | 0.02075 | 0.015881 | 56 | 1.306609 | 0.892558 |
| High Today - High Future | 0.028375 | 0.015881 | 56 | 1.786749 | 0.631361 |
| High Today - High eT | -0.00413 | 0.015881 | 56 | -0.25975 | 0.999995 |
| High Today - High eCO2 | 0.026375 | 0.015881 | 56 | 1.660811 | 0.711623 |
| High eT - Low Future | 0.03175 | 0.015881 | 56 | 1.99927 | 0.49159 |
| High eT - Low eCO2 | 0.024875 | 0.015881 | 56 | 1.566357 | 0.767734 |
| High eT - High Future | 0.0325 | 0.015881 | 56 | 2.046497 | 0.461224 |
| High eT - High eCO2 | 0.0305 | 0.015881 | 56 | 1.920558 | 0.543139 |
| High eCO2 - Low Future | 0.00125 | 0.015881 | 56 | 0.078711 | 1 |
| High eCO2 - High Future | 0.002 | 0.015881 | 56 | 0.125938 | 1 |
| **Straw Zn concentration** |  |  |  |  |  |
| Low Today - Low Future | -3.1095 | 2.198956 | 56 | -1.41408 | 0.846955 |
| Low Today - Low eT | 0.045 | 2.198956 | 56 | 0.020464 | 1 |
| Low Today - Low eCO2 | 0.134625 | 2.198956 | 56 | 0.061222 | 1 |
| Low Today - High Today | 8.538125 | 2.198956 | 56 | 3.882808 | 0.006268 |
| Low Today - High Future | 7.175 | 2.198956 | 56 | 3.262912 | 0.037086 |
| Low Today - High eT | 6.903 | 2.198956 | 56 | 3.139217 | 0.051172 |
| Low Today - High eCO2 | 8.174625 | 2.198956 | 56 | 3.717502 | 0.010316 |
| Low Future - High Future | 10.2845 | 2.198956 | 56 | 4.676992 | 0.000474 |
| Low eT - Low Future | -3.1545 | 2.198956 | 56 | -1.43454 | 0.837252 |
| Low eT - Low eCO2 | 0.089625 | 2.198956 | 56 | 0.040758 | 1 |
| Low eT - High Future | 7.13 | 2.198956 | 56 | 3.242447 | 0.039148 |
| Low eT - High eT | 6.858 | 2.198956 | 56 | 3.118752 | 0.053908 |
| Low eT - High eCO2 | 8.129625 | 2.198956 | 56 | 3.697038 | 0.01096 |
| Low eCO2 - Low Future | -3.24413 | 2.198956 | 56 | -1.4753 | 0.817012 |
| Low eCO2 - High Future | 7.040375 | 2.198956 | 56 | 3.201689 | 0.043559 |
| Low eCO2 - High eCO2 | 8.04 | 2.198956 | 56 | 3.65628 | 0.012355 |
| High Today - Low Future | -11.6476 | 2.198956 | 56 | -5.29689 | 5.37E-05 |
| High Today - Low eT | -8.49313 | 2.198956 | 56 | -3.86234 | 0.006673 |
| High Today - Low eCO2 | -8.4035 | 2.198956 | 56 | -3.82159 | 0.007552 |
| High Today - High Future | -1.36313 | 2.198956 | 56 | -0.6199 | 0.998455 |
| High Today - High eT | -1.63513 | 2.198956 | 56 | -0.74359 | 0.99516 |
| High Today - High eCO2 | -0.3635 | 2.198956 | 56 | -0.16531 | 1 |
| High eT - Low Future | -10.0125 | 2.198956 | 56 | -4.5533 | 0.000721 |
| High eT - Low eCO2 | -6.76838 | 2.198956 | 56 | -3.07799 | 0.059736 |
| High eT - High Future | 0.272 | 2.198956 | 56 | 0.123695 | 1 |
| High eT - High eCO2 | 1.271625 | 2.198956 | 56 | 0.578286 | 0.999012 |
| High eCO2 - Low Future | -11.2841 | 2.198956 | 56 | -5.13158 | 9.70E-05 |
| High eCO2 - High Future | -0.99963 | 2.198956 | 56 | -0.45459 | 0.999797 |
| **Straw Cd concentration** |  |  |  |  |  |
| Low Today - Low Future | -8.98238 | 5.662823 | 56 | -1.5862 | 0.756325 |
| Low Today - Low eT | -11.5616 | 5.662823 | 56 | -2.04167 | 0.464301 |
| Low Today - Low eCO2 | 4.6855 | 5.662823 | 56 | 0.827414 | 0.990763 |
| Low Today - High Today | 13.0845 | 5.662823 | 56 | 2.310597 | 0.30634 |
| Low Today - High Future | 4.846 | 5.662823 | 56 | 0.855757 | 0.988726 |
| Low Today - High eT | 0.031125 | 5.662823 | 56 | 0.005496 | 1 |
| Low Today - High eCO2 | 3.08675 | 5.662823 | 56 | 0.54509 | 0.999327 |
| Low Future - High Future | 13.82838 | 5.662823 | 56 | 2.441958 | 0.242113 |
| Low eT - Low Future | 2.57925 | 5.662823 | 56 | 0.455471 | 0.999794 |
| Low eT - Low eCO2 | 16.24713 | 5.662823 | 56 | 2.869086 | 0.098817 |
| Low eT - High Future | 16.40763 | 5.662823 | 56 | 2.897428 | 0.092509 |
| Low eT - High eT | 11.59275 | 5.662823 | 56 | 2.047168 | 0.460797 |
| Low eT - High eCO2 | 14.64838 | 5.662823 | 56 | 2.586762 | 0.182577 |
| Low eCO2 - Low Future | -13.6679 | 5.662823 | 56 | -2.41361 | 0.255157 |
| Low eCO2 - High Future | 0.1605 | 5.662823 | 56 | 0.028343 | 1 |
| Low eCO2 - High eCO2 | -1.59875 | 5.662823 | 56 | -0.28232 | 0.999992 |
| High Today - Low Future | -22.0669 | 5.662823 | 56 | -3.8968 | 0.006005 |
| High Today - Low eT | -24.6461 | 5.662823 | 56 | -4.35227 | 0.001408 |
| High Today - Low eCO2 | -8.399 | 5.662823 | 56 | -1.48318 | 0.812963 |
| High Today - High Future | -8.2385 | 5.662823 | 56 | -1.45484 | 0.827323 |
| High Today - High eT | -13.0534 | 5.662823 | 56 | -2.3051 | 0.309233 |
| High Today - High eCO2 | -9.99775 | 5.662823 | 56 | -1.76551 | 0.645198 |
| High eT - Low Future | -9.0135 | 5.662823 | 56 | -1.5917 | 0.753126 |
| High eT - Low eCO2 | 4.654375 | 5.662823 | 56 | 0.821918 | 0.991123 |
| High eT - High Future | 4.814875 | 5.662823 | 56 | 0.850261 | 0.989146 |
| High eT - High eCO2 | 3.055625 | 5.662823 | 56 | 0.539594 | 0.99937 |
| High eCO2 - Low Future | -12.0691 | 5.662823 | 56 | -2.13129 | 0.408349 |
| High eCO2 - High Future | 1.75925 | 5.662823 | 56 | 0.310667 | 0.999985 |
| **SECTION E – HUSK, PANICLE, AND STRAW ELEMENTAL CONTENTS** | | | |  |  |
| **Husk Fe content** |  |  |  |  |  |
| Low Today - Low Future | 0.0475 | 0.208022 | 55 | 0.228341 | 0.999998 |
| Low Today - Low eT | -0.175 | 0.208022 | 55 | -0.84126 | 0.989791 |
| Low Today - Low eCO2 | -0.39982 | 0.215324 | 55 | -1.85684 | 0.585327 |
| Low Today - High Today | 0.575 | 0.208022 | 55 | 2.764125 | 0.125692 |
| Low Today - High Future | 0.39125 | 0.208022 | 55 | 1.880807 | 0.569494 |
| Low Today - High eT | 0.68375 | 0.208022 | 55 | 3.286905 | 0.035027 |
| Low Today - High eCO2 | 0.26125 | 0.208022 | 55 | 1.255874 | 0.910785 |
| Low Future - High Future | 0.34375 | 0.208022 | 55 | 1.652466 | 0.716731 |
| Low eT - Low Future | 0.2225 | 0.208022 | 55 | 1.069596 | 0.960443 |
| Low eT - Low eCO2 | -0.22482 | 0.215324 | 55 | -1.04411 | 0.96524 |
| Low eT - High Future | 0.56625 | 0.208022 | 55 | 2.722062 | 0.137796 |
| Low eT - High eT | 0.85875 | 0.208022 | 55 | 4.12816 | 0.002959 |
| Low eT - High eCO2 | 0.43625 | 0.208022 | 55 | 2.097129 | 0.429576 |
| Low eCO2 - Low Future | 0.447321 | 0.215324 | 55 | 2.077437 | 0.441862 |
| Low eCO2 - High Future | 0.791071 | 0.215324 | 55 | 3.673871 | 0.011859 |
| Low eCO2 - High eCO2 | 0.661071 | 0.215324 | 55 | 3.070128 | 0.061224 |
| High Today - Low Future | -0.5275 | 0.208022 | 55 | -2.53578 | 0.202562 |
| High Today - Low eT | -0.75 | 0.208022 | 55 | -3.60538 | 0.01447 |
| High Today - Low eCO2 | -0.97482 | 0.215324 | 55 | -4.52724 | 0.000805 |
| High Today - High Future | -0.18375 | 0.208022 | 55 | -0.88332 | 0.986407 |
| High Today - High eT | 0.10875 | 0.208022 | 55 | 0.52278 | 0.999487 |
| High Today - High eCO2 | -0.31375 | 0.208022 | 55 | -1.50825 | 0.799739 |
| High eT - Low Future | -0.63625 | 0.208022 | 55 | -3.05856 | 0.063007 |
| High eT - Low eCO2 | -1.08357 | 0.215324 | 55 | -5.03229 | 0.000142 |
| High eT - High Future | -0.2925 | 0.208022 | 55 | -1.4061 | 0.850581 |
| High eT - High eCO2 | -0.4225 | 0.208022 | 55 | -2.03103 | 0.471286 |
| High eCO2 - Low Future | -0.21375 | 0.208022 | 55 | -1.02753 | 0.968126 |
| High eCO2 - High Future | 0.13 | 0.208022 | 55 | 0.624933 | 0.998369 |
| **Husk Zn content (varIdent)** |  |  |  |  |  |
| Low Today - Low Future | -17.5975 | 17.02437 | 55 | -1.03367 | 0.967079 |
| Low Today - Low eT | -22.9175 | 17.02437 | 55 | -1.34616 | 0.876771 |
| Low Today - Low eCO2 | -41.635 | 17.6219 | 55 | -2.36269 | 0.28007 |
| Low Today - High Today | 54.63625 | 17.02437 | 55 | 3.209296 | 0.042963 |
| Low Today - High Future | 30.3 | 17.02437 | 55 | 1.779802 | 0.635932 |
| Low Today - High eT | 57.48125 | 17.02437 | 55 | 3.37641 | 0.02752 |
| Low Today - High eCO2 | 51.3 | 17.02437 | 55 | 3.013327 | 0.070417 |
| Low Future - High Future | 47.8975 | 17.02437 | 55 | 2.813467 | 0.11259 |
| Low eT - Low Future | 5.32 | 17.02437 | 55 | 0.312493 | 0.999984 |
| Low eT - Low eCO2 | -18.7175 | 17.6219 | 55 | -1.06217 | 0.961886 |
| Low eT - High Future | 53.2175 | 17.02437 | 55 | 3.12596 | 0.053216 |
| Low eT - High eT | 80.39875 | 17.02437 | 55 | 4.722568 | 0.000416 |
| Low eT - High eCO2 | 74.2175 | 17.02437 | 55 | 4.359486 | 0.001403 |
| Low eCO2 - Low Future | 24.0375 | 17.6219 | 55 | 1.36407 | 0.869241 |
| Low eCO2 - High Future | 71.935 | 17.6219 | 55 | 4.082137 | 0.003423 |
| Low eCO2 - High eCO2 | 92.935 | 17.6219 | 55 | 5.273836 | 6.06E-05 |
| High Today - Low Future | -72.2338 | 17.02437 | 55 | -4.24296 | 0.002049 |
| High Today - Low eT | -77.5537 | 17.02437 | 55 | -4.55545 | 0.000732 |
| High Today - Low eCO2 | -96.2712 | 17.6219 | 55 | -5.46316 | 3.07E-05 |
| High Today - High Future | -24.3363 | 17.02437 | 55 | -1.42949 | 0.839605 |
| High Today - High eT | 2.845 | 17.02437 | 55 | 0.167113 | 1 |
| High Today - High eCO2 | -3.33625 | 17.02437 | 55 | -0.19597 | 0.999999 |
| High eT - Low Future | -75.0788 | 17.02437 | 55 | -4.41008 | 0.001188 |
| High eT - Low eCO2 | -99.1162 | 17.6219 | 55 | -5.62461 | 1.71E-05 |
| High eT - High Future | -27.1813 | 17.02437 | 55 | -1.59661 | 0.750216 |
| High eT - High eCO2 | -6.18125 | 17.02437 | 55 | -0.36308 | 0.999955 |
| High eCO2 - Low Future | -68.8975 | 17.02437 | 55 | -4.04699 | 0.003823 |
| High eCO2 - High Future | -21 | 17.02437 | 55 | -1.23353 | 0.918181 |
| **Husk Cd content** |  |  |  |  |  |
| Low Today - Low Future | -14.0554 | 13.00854 | 53 | -1.08047 | 0.95816 |
| Low Today - Low eT | -20.9 | 12.56745 | 53 | -1.66303 | 0.710206 |
| Low Today - Low eCO2 | -5.47536 | 13.00854 | 53 | -0.4209 | 0.999878 |
| Low Today - High Today | 31.38464 | 13.00854 | 53 | 2.412618 | 0.256724 |
| Low Today - High Future | 20.83 | 12.56745 | 53 | 1.657457 | 0.713631 |
| Low Today - High eT | 30.01875 | 12.56745 | 53 | 2.388612 | 0.268119 |
| Low Today - High eCO2 | 0.375 | 12.56745 | 53 | 0.029839 | 1 |
| Low Future - High Future | 34.88536 | 13.00854 | 53 | 2.681727 | 0.15105 |
| Low eT - Low Future | 6.844643 | 13.00854 | 53 | 0.526165 | 0.999462 |
| Low eT - Low eCO2 | 15.42464 | 13.00854 | 53 | 1.185732 | 0.932517 |
| Low eT - High Future | 41.73 | 12.56745 | 53 | 3.320484 | 0.032493 |
| Low eT - High eT | 50.91875 | 12.56745 | 53 | 4.051638 | 0.003891 |
| Low eT - High eCO2 | 21.275 | 12.56745 | 53 | 1.692866 | 0.691654 |
| Low eCO2 - Low Future | -8.58 | 13.43517 | 53 | -0.63862 | 0.99812 |
| Low eCO2 - High Future | 26.30536 | 13.00854 | 53 | 2.02216 | 0.477343 |
| Low eCO2 - High eCO2 | 5.850357 | 13.00854 | 53 | 0.449732 | 0.99981 |
| High Today - Low Future | -45.44 | 13.43517 | 53 | -3.38217 | 0.027524 |
| High Today - Low eT | -52.2846 | 13.00854 | 53 | -4.01925 | 0.004302 |
| High Today - Low eCO2 | -36.86 | 13.43517 | 53 | -2.74355 | 0.132298 |
| High Today - High Future | -10.5546 | 13.00854 | 53 | -0.81136 | 0.991739 |
| High Today - High eT | -1.36589 | 13.00854 | 53 | -0.105 | 1 |
| High Today - High eCO2 | -31.0096 | 13.00854 | 53 | -2.38379 | 0.270446 |
| High eT - Low Future | -44.0741 | 13.00854 | 53 | -3.38809 | 0.027086 |
| High eT - Low eCO2 | -35.4941 | 13.00854 | 53 | -2.72852 | 0.136677 |
| High eT - High Future | -9.18875 | 12.56745 | 53 | -0.73115 | 0.995612 |
| High eT - High eCO2 | -29.6438 | 12.56745 | 53 | -2.35877 | 0.282728 |
| High eCO2 - Low Future | -14.4304 | 13.00854 | 53 | -1.1093 | 0.951959 |
| High eCO2 - High Future | 20.455 | 12.56745 | 53 | 1.627618 | 0.731748 |
| **Panicle Fe content** |  |  |  |  |  |
| Low Today - Low Future | -10.4834 | 5.172185 | 26.9088 | -2.02688 | 0.48369 |
| Low Today - Low eT | -3.54714 | 5.172185 | 26.9088 | -0.68581 | 0.996726 |
| Low Today - Low eCO2 | -11.8171 | 5.172185 | 26.9088 | -2.28475 | 0.337482 |
| Low Today - High Today | 7.854524 | 5.117335 | 52.73347 | 1.534885 | 0.785174 |
| Low Today - High Future | 6.909107 | 4.817386 | 50.73065 | 1.434203 | 0.837019 |
| Low Today - High eT | 13.05911 | 4.817386 | 50.73065 | 2.710828 | 0.142961 |
| Low Today - High eCO2 | 2.831607 | 4.817386 | 50.73065 | 0.587789 | 0.998888 |
| Low Future - High Future | 17.3925 | 4.628582 | 51.87686 | 3.75763 | 0.009626 |
| Low eT - Low Future | -6.93625 | 4.996806 | 26.9088 | -1.38814 | 0.854699 |
| Low eT - Low eCO2 | -8.27 | 4.996806 | 26.9088 | -1.65506 | 0.714178 |
| Low eT - High Future | 10.45625 | 4.628582 | 51.87686 | 2.259061 | 0.335385 |
| Low eT - High eT | 16.60625 | 4.628582 | 51.87686 | 3.587762 | 0.015721 |
| Low eT - High eCO2 | 6.37875 | 4.628582 | 51.87686 | 1.378122 | 0.86291 |
| Low eCO2 - Low Future | 1.33375 | 4.996806 | 26.9088 | 0.266921 | 0.999994 |
| Low eCO2 - High Future | 18.72625 | 4.628582 | 51.87686 | 4.045785 | 0.004038 |
| Low eCO2 - High eCO2 | 14.64875 | 4.628582 | 51.87686 | 3.164846 | 0.049115 |
| High Today - Low Future | -18.3379 | 4.940009 | 52.9989 | -3.71212 | 0.010847 |
| High Today - Low eT | -11.4017 | 4.940009 | 52.9989 | -2.30803 | 0.308685 |
| High Today - Low eCO2 | -19.6717 | 4.940009 | 52.9989 | -3.98211 | 0.004824 |
| High Today - High Future | -0.94542 | 4.567209 | 25.91078 | -0.207 | 0.999999 |
| High Today - High eT | 5.204583 | 4.567209 | 25.91078 | 1.139555 | 0.94151 |
| High Today - High eCO2 | -5.02292 | 4.567209 | 25.91078 | -1.09978 | 0.951102 |
| High eT - Low Future | -23.5425 | 4.628582 | 51.87686 | -5.08633 | 0.000132 |
| High eT - Low eCO2 | -24.8763 | 4.628582 | 51.87686 | -5.37449 | 4.83E-05 |
| High eT - High Future | -6.15 | 4.228414 | 25.91078 | -1.45445 | 0.823466 |
| High eT - High eCO2 | -10.2275 | 4.228414 | 25.91078 | -2.41876 | 0.27415 |
| High eCO2 - Low Future | -13.315 | 4.628582 | 51.87686 | -2.87669 | 0.098636 |
| High eCO2 - High Future | 4.0775 | 4.228414 | 25.91078 | 0.96431 | 0.975632 |
| **Panicle Zn content (varIdent)** |  |  |  |  |  |
| Low Today - Low Future | -12.3488 | 4.921667 | 27.00099 | -2.50906 | 0.23402 |
| Low Today - Low eT | -4.3775 | 4.921667 | 27.00099 | -0.88943 | 0.984591 |
| Low Today - Low eCO2 | -7.3275 | 4.921667 | 27.00099 | -1.48882 | 0.806544 |
| Low Today - High Today | 3.1075 | 3.778042 | 32.61503 | 0.822516 | 0.990493 |
| Low Today - High Future | 0.62875 | 3.778042 | 32.61503 | 0.166422 | 1 |
| Low Today - High eT | 3.07625 | 3.778042 | 32.61503 | 0.814245 | 0.991044 |
| Low Today - High eCO2 | 1.3425 | 3.778042 | 32.61503 | 0.355343 | 0.999958 |
| Low Future - High Future | 12.9775 | 3.557914 | 33.39819 | 3.647503 | 0.017956 |
| Low eT - Low Future | -7.97125 | 4.754782 | 27.00099 | -1.67647 | 0.701408 |
| Low eT - Low eCO2 | -2.95 | 4.754782 | 27.00099 | -0.62043 | 0.998253 |
| Low eT - High Future | 5.00625 | 3.557914 | 33.39819 | 1.407075 | 0.84762 |
| Low eT - High eT | 7.45375 | 3.557914 | 33.39819 | 2.094978 | 0.43844 |
| Low eT - High eCO2 | 5.72 | 3.557914 | 33.39819 | 1.607684 | 0.742493 |
| Low eCO2 - Low Future | -5.02125 | 4.754782 | 27.00099 | -1.05604 | 0.960548 |
| Low eCO2 - High Future | 7.95625 | 3.557914 | 33.39819 | 2.236212 | 0.357178 |
| Low eCO2 - High eCO2 | 8.67 | 3.557914 | 33.39819 | 2.436821 | 0.25739 |
| High Today - Low Future | -15.4563 | 3.557914 | 33.39819 | -4.34419 | 0.002781 |
| High Today - Low eT | -7.485 | 3.557914 | 33.39819 | -2.10376 | 0.433168 |
| High Today - Low eCO2 | -10.435 | 3.557914 | 33.39819 | -2.9329 | 0.097667 |
| High Today - High Future | -2.47875 | 1.64607 | 27.99989 | -1.50586 | 0.797961 |
| High Today - High eT | -0.03125 | 1.64607 | 27.99989 | -0.01898 | 1 |
| High Today - High eCO2 | -1.765 | 1.64607 | 27.99989 | -1.07225 | 0.957468 |
| High eT - Low Future | -15.425 | 3.557914 | 33.39819 | -4.33541 | 0.00285 |
| High eT - Low eCO2 | -10.4038 | 3.557914 | 33.39819 | -2.92412 | 0.099527 |
| High eT - High Future | -2.4475 | 1.64607 | 27.99989 | -1.48687 | 0.807767 |
| High eT - High eCO2 | -1.73375 | 1.64607 | 27.99989 | -1.05327 | 0.961258 |
| High eCO2 - Low Future | -13.6913 | 3.557914 | 33.39819 | -3.84811 | 0.010671 |
| High eCO2 - High Future | -0.71375 | 1.64607 | 27.99989 | -0.43361 | 0.999832 |
| **Panicle Cd content** |  |  |  |  |  |
| Low Today - Low Future | -8.34625 | 3.293963 | 55 | -2.5338 | 0.203354 |
| Low Today - Low eT | -8.99125 | 3.293963 | 55 | -2.72961 | 0.135558 |
| Low Today - Low eCO2 | -4.76875 | 3.293963 | 55 | -1.44772 | 0.83077 |
| Low Today - High Today | -0.0825 | 3.293963 | 55 | -0.02505 | 1 |
| Low Today - High Future | -3.8575 | 3.293963 | 55 | -1.17108 | 0.936718 |
| Low Today - High eT | -1.255 | 3.293963 | 55 | -0.381 | 0.999938 |
| Low Today - High eCO2 | -4.0625 | 3.293963 | 55 | -1.23332 | 0.918248 |
| Low Future - High Future | 4.48875 | 3.182271 | 55 | 1.410549 | 0.848525 |
| Low eT - Low Future | 0.645 | 3.182271 | 55 | 0.202685 | 0.999999 |
| Low eT - Low eCO2 | 4.2225 | 3.182271 | 55 | 1.326883 | 0.884588 |
| Low eT - High Future | 5.13375 | 3.182271 | 55 | 1.613235 | 0.740406 |
| Low eT - High eT | 7.73625 | 3.182271 | 55 | 2.431047 | 0.247439 |
| Low eT - High eCO2 | 4.92875 | 3.182271 | 55 | 1.548815 | 0.77758 |
| Low eCO2 - Low Future | -3.5775 | 3.182271 | 55 | -1.1242 | 0.94863 |
| Low eCO2 - High Future | 0.91125 | 3.182271 | 55 | 0.286352 | 0.999991 |
| Low eCO2 - High eCO2 | 0.70625 | 3.182271 | 55 | 0.221933 | 0.999998 |
| High Today - Low Future | -8.26375 | 3.182271 | 55 | -2.59681 | 0.179266 |
| High Today - Low eT | -8.90875 | 3.182271 | 55 | -2.79949 | 0.116183 |
| High Today - Low eCO2 | -4.68625 | 3.182271 | 55 | -1.47261 | 0.818319 |
| High Today - High Future | -3.775 | 3.182271 | 55 | -1.18626 | 0.932497 |
| High Today - High eT | -1.1725 | 3.182271 | 55 | -0.36845 | 0.99995 |
| High Today - High eCO2 | -3.98 | 3.182271 | 55 | -1.25068 | 0.91254 |
| High eT - Low Future | -7.09125 | 3.182271 | 55 | -2.22836 | 0.351527 |
| High eT - Low eCO2 | -3.51375 | 3.182271 | 55 | -1.10416 | 0.953215 |
| High eT - High Future | -2.6025 | 3.182271 | 55 | -0.81781 | 0.99137 |
| High eT - High eCO2 | -2.8075 | 3.182271 | 55 | -0.88223 | 0.986504 |
| High eCO2 - Low Future | -4.28375 | 3.182271 | 55 | -1.34613 | 0.876783 |
| High eCO2 - High Future | 0.205 | 3.182271 | 55 | 0.064419 | 1 |
| **Straw Fe content** |  |  |  |  |  |
| Low Today - Low Future | 0.060536 | 0.183941 | 55 | 0.329105 | 0.999977 |
| Low Today - Low eT | 0.090536 | 0.183941 | 55 | 0.492201 | 0.999655 |
| Low Today - Low eCO2 | 0.063036 | 0.183941 | 55 | 0.342696 | 0.99997 |
| Low Today - High Today | 0.554286 | 0.183941 | 55 | 3.013394 | 0.070406 |
| Low Today - High Future | 0.551786 | 0.183941 | 55 | 2.999802 | 0.072773 |
| Low Today - High eT | 0.396786 | 0.183941 | 55 | 2.15714 | 0.392998 |
| Low Today - High eCO2 | 0.381786 | 0.183941 | 55 | 2.075591 | 0.44302 |
| Low Future - High Future | 0.49125 | 0.177704 | 55 | 2.764435 | 0.125606 |
| Low eT - Low Future | -0.03 | 0.177704 | 55 | -0.16882 | 1 |
| Low eT - Low eCO2 | -0.0275 | 0.177704 | 55 | -0.15475 | 1 |
| Low eT - High Future | 0.46125 | 0.177704 | 55 | 2.595615 | 0.179702 |
| Low eT - High eT | 0.30625 | 0.177704 | 55 | 1.723375 | 0.672345 |
| Low eT - High eCO2 | 0.29125 | 0.177704 | 55 | 1.638965 | 0.724958 |
| Low eCO2 - Low Future | -0.0025 | 0.177704 | 55 | -0.01407 | 1 |
| Low eCO2 - High Future | 0.48875 | 0.177704 | 55 | 2.750367 | 0.129555 |
| Low eCO2 - High eCO2 | 0.31875 | 0.177704 | 55 | 1.793717 | 0.626847 |
| High Today - Low Future | -0.49375 | 0.177704 | 55 | -2.7785 | 0.121754 |
| High Today - Low eT | -0.46375 | 0.177704 | 55 | -2.60968 | 0.174615 |
| High Today - Low eCO2 | -0.49125 | 0.177704 | 55 | -2.76443 | 0.125606 |
| High Today - High Future | -0.0025 | 0.177704 | 55 | -0.01407 | 1 |
| High Today - High eT | -0.1575 | 0.177704 | 55 | -0.88631 | 0.986137 |
| High Today - High eCO2 | -0.1725 | 0.177704 | 55 | -0.97072 | 0.976712 |
| High eT - Low Future | -0.33625 | 0.177704 | 55 | -1.8922 | 0.56197 |
| High eT - Low eCO2 | -0.33375 | 0.177704 | 55 | -1.87813 | 0.571264 |
| High eT - High Future | 0.155 | 0.177704 | 55 | 0.872239 | 0.987371 |
| High eT - High eCO2 | -0.015 | 0.177704 | 55 | -0.08441 | 1 |
| High eCO2 - Low Future | -0.32125 | 0.177704 | 55 | -1.80779 | 0.617631 |
| High eCO2 - High Future | 0.17 | 0.177704 | 55 | 0.956649 | 0.978543 |
| **Straw Zn content** |  |  |  |  |  |
| Low Today - Low Future | -111.797 | 27.0923 | 55 | -4.12651 | 0.002975 |
| Low Today - Low eT | -38.243 | 27.0923 | 55 | -1.41158 | 0.848045 |
| Low Today - Low eCO2 | -51.4318 | 27.0923 | 55 | -1.89839 | 0.557878 |
| Low Today - High Today | 93.59946 | 27.0923 | 55 | 3.454836 | 0.022171 |
| Low Today - High Future | 67.71571 | 27.0923 | 55 | 2.499445 | 0.217428 |
| Low Today - High eT | 71.41071 | 27.0923 | 55 | 2.63583 | 0.165448 |
| Low Today - High eCO2 | 61.38696 | 27.0923 | 55 | 2.265845 | 0.330653 |
| Low Future - High Future | 179.5125 | 26.17365 | 55 | 6.85852 | 1.76E-07 |
| Low eT - Low Future | -73.5538 | 26.17365 | 55 | -2.81022 | 0.113417 |
| Low eT - Low eCO2 | -13.1888 | 26.17365 | 55 | -0.50389 | 0.999597 |
| Low eT - High Future | 105.9588 | 26.17365 | 55 | 4.048299 | 0.003807 |
| Low eT - High eT | 109.6538 | 26.17365 | 55 | 4.189471 | 0.002434 |
| Low eT - High eCO2 | 99.63 | 26.17365 | 55 | 3.8065 | 0.008002 |
| Low eCO2 - Low Future | -60.365 | 26.17365 | 55 | -2.30633 | 0.308905 |
| Low eCO2 - High Future | 119.1475 | 26.17365 | 55 | 4.552193 | 0.00074 |
| Low eCO2 - High eCO2 | 112.8188 | 26.17365 | 55 | 4.310394 | 0.001647 |
| High Today - Low Future | -205.396 | 26.17365 | 55 | -7.84744 | 4.28E-09 |
| High Today - Low eT | -131.843 | 26.17365 | 55 | -5.03722 | 0.00014 |
| High Today - Low eCO2 | -145.031 | 26.17365 | 55 | -5.54112 | 2.32E-05 |
| High Today - High Future | -25.8837 | 26.17365 | 55 | -0.98892 | 0.974173 |
| High Today - High eT | -22.1887 | 26.17365 | 55 | -0.84775 | 0.989316 |
| High Today - High eCO2 | -32.2125 | 26.17365 | 55 | -1.23072 | 0.91908 |
| High eT - Low Future | -183.208 | 26.17365 | 55 | -6.99969 | 1.04E-07 |
| High eT - Low eCO2 | -122.843 | 26.17365 | 55 | -4.69337 | 0.00046 |
| High eT - High Future | -3.695 | 26.17365 | 55 | -0.14117 | 1 |
| High eT - High eCO2 | -10.0238 | 26.17365 | 55 | -0.38297 | 0.999936 |
| High eCO2 - Low Future | -173.184 | 26.17365 | 55 | -6.61672 | 4.36E-07 |
| High eCO2 - High Future | 6.32875 | 26.17365 | 55 | 0.241799 | 0.999997 |
| **Straw Cd content** |  |  |  |  |  |
| Low Today - Low Future | -175.69 | 70.24387 | 56 | -2.50114 | 0.216344 |
| Low Today - Low eT | -157.268 | 70.24387 | 56 | -2.23888 | 0.345311 |
| Low Today - Low eCO2 | 6.27875 | 70.24387 | 56 | 0.089385 | 1 |
| Low Today - High Today | 191.0113 | 70.24387 | 56 | 2.719259 | 0.138256 |
| Low Today - High Future | 95.02 | 70.24387 | 56 | 1.352716 | 0.874118 |
| Low Today - High eT | 67.09125 | 70.24387 | 56 | 0.955119 | 0.978765 |
| Low Today - High eCO2 | 37.22875 | 70.24387 | 56 | 0.529993 | 0.99944 |
| Low Future - High Future | 270.71 | 70.24387 | 56 | 3.853859 | 0.006847 |
| Low eT - Low Future | -18.4225 | 70.24387 | 56 | -0.26226 | 0.999995 |
| Low eT - Low eCO2 | 163.5463 | 70.24387 | 56 | 2.328264 | 0.29715 |
| Low eT - High Future | 252.2875 | 70.24387 | 56 | 3.591594 | 0.014911 |
| Low eT - High eT | 224.3588 | 70.24387 | 56 | 3.193998 | 0.044439 |
| Low eT - High eCO2 | 194.4963 | 70.24387 | 56 | 2.768871 | 0.124008 |
| Low eCO2 - Low Future | -181.969 | 70.24387 | 56 | -2.59053 | 0.181186 |
| Low eCO2 - High Future | 88.74125 | 70.24387 | 56 | 1.263331 | 0.908296 |
| Low eCO2 - High eCO2 | 30.95 | 70.24387 | 56 | 0.440608 | 0.999835 |
| High Today - Low Future | -366.701 | 70.24387 | 56 | -5.2204 | 7.07E-05 |
| High Today - Low eT | -348.279 | 70.24387 | 56 | -4.95814 | 0.000179 |
| High Today - Low eCO2 | -184.733 | 70.24387 | 56 | -2.62987 | 0.167121 |
| High Today - High Future | -95.9913 | 70.24387 | 56 | -1.36654 | 0.868255 |
| High Today - High eT | -123.92 | 70.24387 | 56 | -1.76414 | 0.646085 |
| High Today - High eCO2 | -153.783 | 70.24387 | 56 | -2.18927 | 0.373745 |
| High eT - Low Future | -242.781 | 70.24387 | 56 | -3.45626 | 0.0219 |
| High eT - Low eCO2 | -60.8125 | 70.24387 | 56 | -0.86573 | 0.987931 |
| High eT - High Future | 27.92875 | 70.24387 | 56 | 0.397597 | 0.999917 |
| High eT - High eCO2 | -29.8625 | 70.24387 | 56 | -0.42513 | 0.99987 |
| High eCO2 - Low Future | -212.919 | 70.24387 | 56 | -3.03114 | 0.067101 |
| High eCO2 - High Future | 57.79125 | 70.24387 | 56 | 0.822723 | 0.991071 |
| **Straw Fe retention factor** |  |  |  |  |  |
| Low Today - Low Future | 0.035714 | 0.060124 | 53 | 0.594014 | 0.998817 |
| Low Today - Low eT | 0.068214 | 0.060124 | 53 | 1.134567 | 0.946023 |
| Low Today - Low eCO2 | 0.051964 | 0.060124 | 53 | 0.86429 | 0.98799 |
| Low Today - High Today | -0.04429 | 0.064631 | 53 | -0.68521 | 0.997068 |
| Low Today - High Future | 0.048214 | 0.060124 | 53 | 0.801919 | 0.992299 |
| Low Today - High eT | -0.12929 | 0.060124 | 53 | -2.15033 | 0.397597 |
| Low Today - High eCO2 | 0.035714 | 0.060124 | 53 | 0.594014 | 0.998817 |
| Low Future - High Future | 0.0125 | 0.058085 | 53 | 0.215202 | 0.999999 |
| Low eT - Low Future | -0.0325 | 0.058085 | 53 | -0.55953 | 0.999197 |
| Low eT - Low eCO2 | -0.01625 | 0.058085 | 53 | -0.27976 | 0.999992 |
| Low eT - High Future | -0.02 | 0.058085 | 53 | -0.34432 | 0.999969 |
| Low eT - High eT | -0.1975 | 0.058085 | 53 | -3.40019 | 0.026208 |
| Low eT - High eCO2 | -0.0325 | 0.058085 | 53 | -0.55953 | 0.999197 |
| Low eCO2 - Low Future | -0.01625 | 0.058085 | 53 | -0.27976 | 0.999992 |
| Low eCO2 - High Future | -0.00375 | 0.058085 | 53 | -0.06456 | 1 |
| Low eCO2 - High eCO2 | -0.01625 | 0.058085 | 53 | -0.27976 | 0.999992 |
| High Today - Low Future | 0.08 | 0.062739 | 53 | 1.275125 | 0.903939 |
| High Today - Low eT | 0.1125 | 0.062739 | 53 | 1.793145 | 0.627311 |
| High Today - Low eCO2 | 0.09625 | 0.062739 | 53 | 1.534135 | 0.785603 |
| High Today - High Future | 0.0925 | 0.062739 | 53 | 1.474364 | 0.817288 |
| High Today - High eT | -0.085 | 0.062739 | 53 | -1.35482 | 0.873007 |
| High Today - High eCO2 | 0.08 | 0.062739 | 53 | 1.275125 | 0.903939 |
| High eT - Low Future | 0.165 | 0.058085 | 53 | 2.840666 | 0.106617 |
| High eT - Low eCO2 | 0.18125 | 0.058085 | 53 | 3.120429 | 0.054574 |
| High eT - High Future | 0.1775 | 0.058085 | 53 | 3.055868 | 0.064078 |
| High eT - High eCO2 | 0.165 | 0.058085 | 53 | 2.840666 | 0.106617 |
| High eCO2 - Low Future | -2.08E-17 | 0.058085 | 53 | -3.58E-16 | 1 |
| High eCO2 - High Future | 0.0125 | 0.058085 | 53 | 0.215202 | 0.999999 |
| **Straw Zn retention factor** |  |  |  |  |  |
| Low Today - Low Future | -0.08393 | 0.051183 | 54 | -1.63977 | 0.724442 |
| Low Today - Low eT | -0.00393 | 0.051183 | 54 | -0.07676 | 1 |
| Low Today - Low eCO2 | -0.01571 | 0.052862 | 54 | -0.29727 | 0.999989 |
| Low Today - High Today | -0.00893 | 0.051183 | 54 | -0.17444 | 1 |
| Low Today - High Future | -0.00268 | 0.051183 | 54 | -0.05233 | 1 |
| Low Today - High eT | -0.15768 | 0.051183 | 54 | -3.08068 | 0.059946 |
| Low Today - High eCO2 | -0.07893 | 0.051183 | 54 | -1.54208 | 0.781273 |
| Low Future - High Future | 0.08125 | 0.049448 | 54 | 1.643157 | 0.722388 |
| Low eT - Low Future | -0.08 | 0.049448 | 54 | -1.61788 | 0.737609 |
| Low eT - Low eCO2 | -0.01179 | 0.051183 | 54 | -0.23027 | 0.999998 |
| Low eT - High Future | 0.00125 | 0.049448 | 54 | 0.025279 | 1 |
| Low eT - High eT | -0.15375 | 0.049448 | 54 | -3.10936 | 0.055798 |
| Low eT - High eCO2 | -0.075 | 0.049448 | 54 | -1.51676 | 0.795117 |
| Low eCO2 - Low Future | -0.06821 | 0.051183 | 54 | -1.33275 | 0.882164 |
| Low eCO2 - High Future | 0.013036 | 0.051183 | 54 | 0.254688 | 0.999996 |
| Low eCO2 - High eCO2 | -0.06321 | 0.051183 | 54 | -1.23506 | 0.917615 |
| High Today - Low Future | -0.075 | 0.049448 | 54 | -1.51676 | 0.795117 |
| High Today - Low eT | 0.005 | 0.049448 | 54 | 0.101117 | 1 |
| High Today - Low eCO2 | -0.00679 | 0.051183 | 54 | -0.13258 | 1 |
| High Today - High Future | 0.00625 | 0.049448 | 54 | 0.126397 | 1 |
| High Today - High eT | -0.14875 | 0.049448 | 54 | -3.00824 | 0.071628 |
| High Today - High eCO2 | -0.07 | 0.049448 | 54 | -1.41564 | 0.846079 |
| High eT - Low Future | 0.07375 | 0.049448 | 54 | 1.491481 | 0.808524 |
| High eT - Low eCO2 | 0.141964 | 0.051183 | 54 | 2.773659 | 0.123455 |
| High eT - High Future | 0.155 | 0.049448 | 54 | 3.134637 | 0.05235 |
| High eT - High eCO2 | 0.07875 | 0.049448 | 54 | 1.592598 | 0.752519 |
| High eCO2 - Low Future | -0.005 | 0.049448 | 54 | -0.10112 | 1 |
| High eCO2 - High Future | 0.07625 | 0.049448 | 54 | 1.542039 | 0.781298 |
| **Straw Cd retention factor** |  |  |  |  |  |
| Low Today - Low Future | 0.185 | 0.051781 | 24 | 3.572754 | 0.028118 |
| Low Today - Low eT | 0.165 | 0.051781 | 24 | 3.18651 | 0.065318 |
| Low Today - Low eCO2 | 0.0175 | 0.051781 | 24 | 0.337963 | 0.999967 |
| Low Today - High Today | -0.19 | 0.051781 | 24 | -3.66931 | 0.022591 |
| Low Today - High Future | 0.0025 | 0.051781 | 24 | 0.04828 | 1 |
| Low Today - High eT | -0.0575 | 0.051781 | 24 | -1.11045 | 0.94818 |
| Low Today - High eCO2 | 0.0275 | 0.051781 | 24 | 0.531085 | 0.999337 |
| Low Future - High Future | -0.1825 | 0.051781 | 24 | -3.52447 | 0.031336 |
| Low eT - Low Future | 0.02 | 0.051781 | 24 | 0.386244 | 0.99992 |
| Low eT - Low eCO2 | -0.1475 | 0.051781 | 24 | -2.84855 | 0.129176 |
| Low eT - High Future | -0.1625 | 0.051781 | 24 | -3.13823 | 0.072261 |
| Low eT - High eT | -0.2225 | 0.051781 | 24 | -4.29696 | 0.005164 |
| Low eT - High eCO2 | -0.1375 | 0.051781 | 24 | -2.65543 | 0.185041 |
| Low eCO2 - Low Future | 0.1675 | 0.051781 | 24 | 3.23479 | 0.058979 |
| Low eCO2 - High Future | -0.015 | 0.051781 | 24 | -0.28968 | 0.999989 |
| Low eCO2 - High eCO2 | 0.01 | 0.051781 | 24 | 0.193122 | 0.999999 |
| High Today - Low Future | 0.375 | 0.051781 | 24 | 7.242068 | 4.37E-06 |
| High Today - Low eT | 0.355 | 0.051781 | 24 | 6.855825 | 1.07E-05 |
| High Today - Low eCO2 | 0.2075 | 0.051781 | 24 | 4.007278 | 0.010303 |
| High Today - High Future | 0.1925 | 0.051781 | 24 | 3.717595 | 0.020229 |
| High Today - High eT | 0.1325 | 0.051781 | 24 | 2.558864 | 0.219327 |
| High Today - High eCO2 | 0.2175 | 0.051781 | 24 | 4.2004 | 0.00651 |
| High eT - Low Future | 0.2425 | 0.051781 | 24 | 4.683204 | 0.002024 |
| High eT - Low eCO2 | 0.075 | 0.051781 | 24 | 1.448414 | 0.825843 |
| High eT - High Future | 0.06 | 0.051781 | 24 | 1.158731 | 0.935954 |
| High eT - High eCO2 | 0.085 | 0.051781 | 24 | 1.641535 | 0.721913 |
| High eCO2 - Low Future | 0.1575 | 0.051781 | 24 | 3.041669 | 0.088139 |
| High eCO2 - High Future | -0.025 | 0.051781 | 24 | -0.4828 | 0.999645 |
| **Husk Fe retention factor** |  |  |  |  |  |
| Low Today - Low Future | -0.01375 | 0.027142 | 55 | -0.50659 | 0.999583 |
| Low Today - Low eT | -0.025 | 0.027142 | 55 | -0.92108 | 0.982695 |
| Low Today - Low eCO2 | -0.04732 | 0.028095 | 55 | -1.68435 | 0.697 |
| Low Today - High Today | 0.01375 | 0.027142 | 55 | 0.506592 | 0.999583 |
| Low Today - High Future | -0.0225 | 0.027142 | 55 | -0.82897 | 0.990644 |
| Low Today - High eT | -0.0225 | 0.027142 | 55 | -0.82897 | 0.990644 |
| Low Today - High eCO2 | -0.0325 | 0.027142 | 55 | -1.1974 | 0.929284 |
| Low Future - High Future | -0.00875 | 0.027142 | 55 | -0.32238 | 0.99998 |
| Low eT - Low Future | 0.01125 | 0.027142 | 55 | 0.414484 | 0.99989 |
| Low eT - Low eCO2 | -0.02232 | 0.028095 | 55 | -0.7945 | 0.992744 |
| Low eT - High Future | 0.0025 | 0.027142 | 55 | 0.092108 | 1 |
| Low eT - High eT | 0.0025 | 0.027142 | 55 | 0.092108 | 1 |
| Low eT - High eCO2 | -0.0075 | 0.027142 | 55 | -0.27632 | 0.999993 |
| Low eCO2 - Low Future | 0.033571 | 0.028095 | 55 | 1.194934 | 0.930004 |
| Low eCO2 - High Future | 0.024821 | 0.028095 | 55 | 0.883488 | 0.986392 |
| Low eCO2 - High eCO2 | 0.014821 | 0.028095 | 55 | 0.527551 | 0.999455 |
| High Today - Low Future | -0.0275 | 0.027142 | 55 | -1.01318 | 0.970482 |
| High Today - Low eT | -0.03875 | 0.027142 | 55 | -1.42767 | 0.840476 |
| High Today - Low eCO2 | -0.06107 | 0.028095 | 55 | -2.17376 | 0.383118 |
| High Today - High Future | -0.03625 | 0.027142 | 55 | -1.33556 | 0.881106 |
| High Today - High eT | -0.03625 | 0.027142 | 55 | -1.33556 | 0.881106 |
| High Today - High eCO2 | -0.04625 | 0.027142 | 55 | -1.70399 | 0.684654 |
| High eT - Low Future | 0.00875 | 0.027142 | 55 | 0.322377 | 0.99998 |
| High eT - Low eCO2 | -0.02482 | 0.028095 | 55 | -0.88349 | 0.986392 |
| High eT - High Future | -7.46E-17 | 0.027142 | 55 | -2.75E-15 | 1 |
| High eT - High eCO2 | -0.01 | 0.027142 | 55 | -0.36843 | 0.999951 |
| High eCO2 - Low Future | 0.01875 | 0.027142 | 55 | 0.690807 | 0.996928 |
| High eCO2 - High Future | 0.01 | 0.027142 | 55 | 0.368431 | 0.999951 |
| **Husk Zn retention factor (varIdent)** |  |  |  |  |  |
| Low Today - Low Future | -0.01875 | 0.026266 | 27.00485 | -0.71384 | 0.995813 |
| Low Today - Low eT | -0.025 | 0.026266 | 27.00485 | -0.95179 | 0.97746 |
| Low Today - Low eCO2 | -0.05268 | 0.027188 | 27.00485 | -1.93756 | 0.538874 |
| Low Today - High Today | -0.01375 | 0.042221 | 40.62563 | -0.32567 | 0.999978 |
| Low Today - High Future | -0.02375 | 0.042221 | 40.62563 | -0.56251 | 0.999136 |
| Low Today - High eT | -0.03875 | 0.042221 | 40.62563 | -0.91779 | 0.982549 |
| Low Today - High eCO2 | -0.035 | 0.042221 | 40.62563 | -0.82897 | 0.990335 |
| Low Future - High Future | -0.005 | 0.042221 | 40.62563 | -0.11842 | 1 |
| Low eT - Low Future | 0.00625 | 0.026266 | 27.00485 | 0.237948 | 0.999997 |
| Low eT - Low eCO2 | -0.02768 | 0.027188 | 27.00485 | -1.01804 | 0.967502 |
| Low eT - High Future | 0.00125 | 0.042221 | 40.62563 | 0.029606 | 1 |
| Low eT - High eT | -0.01375 | 0.042221 | 40.62563 | -0.32567 | 0.999978 |
| Low eT - High eCO2 | -0.01 | 0.042221 | 40.62563 | -0.23685 | 0.999997 |
| Low eCO2 - Low Future | 0.033929 | 0.027188 | 27.00485 | 1.247919 | 0.909651 |
| Low eCO2 - High Future | 0.028929 | 0.042801 | 42.17509 | 0.675889 | 0.997231 |
| Low eCO2 - High eCO2 | 0.017679 | 0.042801 | 42.17509 | 0.413044 | 0.999889 |
| High Today - Low Future | -0.005 | 0.042221 | 40.62563 | -0.11842 | 1 |
| High Today - Low eT | -0.01125 | 0.042221 | 40.62563 | -0.26645 | 0.999994 |
| High Today - Low eCO2 | -0.03893 | 0.042801 | 42.17509 | -0.90953 | 0.983492 |
| High Today - High Future | -0.01 | 0.053622 | 28.00338 | -0.18649 | 0.999999 |
| High Today - High eT | -0.025 | 0.053622 | 28.00338 | -0.46623 | 0.999728 |
| High Today - High eCO2 | -0.02125 | 0.053622 | 28.00338 | -0.39629 | 0.999908 |
| High eT - Low Future | 0.02 | 0.042221 | 40.62563 | 0.473696 | 0.99972 |
| High eT - Low eCO2 | -0.01393 | 0.042801 | 42.17509 | -0.32543 | 0.999978 |
| High eT - High Future | 0.015 | 0.053622 | 28.00338 | 0.279735 | 0.999991 |
| High eT - High eCO2 | 0.00375 | 0.053622 | 28.00338 | 0.069934 | 1 |
| High eCO2 - Low Future | 0.01625 | 0.042221 | 40.62563 | 0.384878 | 0.99993 |
| High eCO2 - High Future | 0.01125 | 0.053622 | 28.00338 | 0.209801 | 0.999999 |
| **Husk Cd retention factor (varIdent)** |  |  |  |  |  |
| Low Today - Low Future | 0.0675 | 0.080124 | 12.01477 | 0.842448 | 0.986353 |
| Low Today - Low eT | 0.155 | 0.080124 | 12.01477 | 1.934511 | 0.555012 |
| Low Today - Low eCO2 | -0.035 | 0.080124 | 12.01477 | -0.43683 | 0.999764 |
| Low Today - High Today | -0.1125 | 0.080497 | 23.99794 | -1.39758 | 0.849591 |
| Low Today - High Future | 0.085 | 0.080497 | 23.99794 | 1.055946 | 0.959968 |
| Low Today - High eT | 0.135 | 0.080497 | 23.99794 | 1.677091 | 0.700952 |
| Low Today - High eCO2 | -0.0925 | 0.080497 | 23.99794 | -1.14912 | 0.938525 |
| Low Future - High Future | 0.0175 | 0.080497 | 23.99794 | 0.217401 | 0.999998 |
| Low eT - Low Future | -0.0875 | 0.080124 | 12.01477 | -1.09206 | 0.946776 |
| Low eT - Low eCO2 | -0.19 | 0.080124 | 12.01477 | -2.37134 | 0.332945 |
| Low eT - High Future | -0.07 | 0.080497 | 23.99794 | -0.8696 | 0.986179 |
| Low eT - High eT | -0.02 | 0.080497 | 23.99794 | -0.24846 | 0.999996 |
| Low eT - High eCO2 | -0.2475 | 0.080497 | 23.99794 | -3.07467 | 0.082401 |
| Low eCO2 - Low Future | 0.1025 | 0.080124 | 12.01477 | 1.279273 | 0.890312 |
| Low eCO2 - High Future | 0.12 | 0.080497 | 23.99794 | 1.490748 | 0.80479 |
| Low eCO2 - High eCO2 | -0.0575 | 0.080497 | 23.99794 | -0.71432 | 0.995692 |
| High Today - Low Future | 0.18 | 0.080497 | 23.99794 | 2.236122 | 0.366575 |
| High Today - Low eT | 0.2675 | 0.080497 | 23.99794 | 3.323126 | 0.048806 |
| High Today - Low eCO2 | 0.0775 | 0.080497 | 23.99794 | 0.962775 | 0.97556 |
| High Today - High Future | 0.1975 | 0.080868 | 12.014 | 2.442261 | 0.303086 |
| High Today - High eT | 0.2475 | 0.080868 | 12.014 | 3.060555 | 0.122457 |
| High Today - High eCO2 | 0.02 | 0.080868 | 12.014 | 0.247318 | 0.999995 |
| High eT - Low Future | -0.0675 | 0.080497 | 23.99794 | -0.83855 | 0.988792 |
| High eT - Low eCO2 | -0.17 | 0.080497 | 23.99794 | -2.11189 | 0.435609 |
| High eT - High Future | -0.05 | 0.080868 | 12.014 | -0.61829 | 0.997812 |
| High eT - High eCO2 | -0.2275 | 0.080868 | 12.014 | -2.81324 | 0.178714 |
| High eCO2 - Low Future | 0.16 | 0.080497 | 23.99794 | 1.987664 | 0.509648 |
| High eCO2 - High Future | 0.1775 | 0.080868 | 12.014 | 2.194944 | 0.415502 |

**Table S4** Grain biomass yield per rice plant

Rice (in Oryza sativa L. cv. M206 grains) was grown on Californian paddy soil bearing either a low (7.3 mg As kg^−1^_dry soil_) or high arsenic (24.5 mg As kg^−1^_dry soil_) content and exposed to four different climatic conditions: today (33ºC, 415 ppmv CO_2_), elevated CO_2_ (↑CO_2_; 33ºC, 850 ppmv CO_2_), elevated temperature (↑T; 38ºC, 415 ppmv CO_2_) and future (38ºC, 850 ppmv CO_2_). Data was previously published in (Muehe et al. 2019). Here, statistical analyses of climate, arsenic, and their interaction on biomass were performed using ANOVA at a 95% confidence level using the results of a generalized least squares test.

| Arsenic | Climatic conditions | Grain yield [g plant^-1^] | | | | | | | p<0.05^#^ |
| --- | --- | --- | --- | --- | --- | --- | --- | --- | --- |
| Low arsenic |  |  | |  | | |  | |  |
|  | Today | | 11.5 | | ± | 2.0 | | ab | |
|  | ↑CO_2_ | | 12.4 | | ± | 2.2 | | a | |
|  | ↑T | | 10.0 | | ± | 1.9 | | abc | |
|  | Future | | 9.6 | | ± | 2.9 | | abc | |
| High arsenic |  | |  | |  |  | |  | |
|  | Today | | 6.9 | | ± | 1.8 | | cd | |
|  | ↑CO_2_ | | 8.7 | | ± | 1.9 | | bc | |
|  | ↑T | | 4.0 | | ± | 2.5 | | d | |
|  | Future | | 6.6 | | ± | 1.3 | | cd | |
| Two Factor ANOVA based on generalized least squares model results^¶‡^ | | | | | | | | | |
|  | dF | Chi squared | |  | | | Pr(>F) | | ^¶^ |
| Arsenic | 1 | 66.236 | |  | | | 4.00E-16 | | *** |
| Climate | 3 | 25.390 | |  | | | 1.28E-05 | | *** |
| Arsenic:Climate | 3 | 4.430 | |  | | | 0.219 | |  |
| Residual | 56 |  | |  | | |  | |  |
| ^#^Different letters indicate differences in grain yields according to Emmeans Post-hoc test  ^¶^Significance codes for two-factor ANOVA analysis: *** < 0.001, ** < 0.01, * < 0.05, ° < 0.1 | | | | | | | | | |

**Table S5** PCA principal components, tissue contributions to principle components, ANOVA testing and model fit statistics.

Rice (in *Oryza sativa* L. cv. M206 grains) was grown on Californian paddy soil bearing either a low (7.3 mg As kg^−1^_dry soil_) or high arsenic (24.5 mg As kg^−1^_dry soil_) content and exposed to four different climatic conditions: today (33ºC, 415 ppmv CO_2_), elevated CO_2_ (33ºC, 850 ppmv CO_2_), elevated (38ºC, 415 ppmv CO_2_), and future (38ºC, 850 ppmv CO_2_). (A) Iron partitioning on low-As soil, (B) Iron partitioning on high-As soil, (C) Zinc partitioning on low-As soil, (D) Zinc partitioning on high-As soil, (E) Cadmium partitioning on low-As soil, (F) Cadmium partitioning on high-As soil. Information on bootstrapping (n = 1000, 95% CI) is provided in Fig. S4.

### A) **IRON partitioning in low-As soil**

#### *Variance explained by principal components:*

| PC | Std_Dev | Proportion_Variance (%) | Cumulative_Variance |
| --- | --- | --- | --- |
| PC1 | 1.775 | 0.4503 | 0.4503 |
| PC2 | 1.416 | 0.2866 | 0.7370 |
| PC3 | 0.980 | 0.1371 | 0.8741 |
| Other PCs below 10% | | | |

#### *Tissue contributions to principal components (squared loadings, %):*

| Variable | PC1 | PC2 | PC3 |
| --- | --- | --- | --- |
| gFeKgStraw | 0.105 | 0.248 | 0.022 |
| mgFeKgPanicle | 0.053 | 0.011 | 0.735 |
| gFeKgHusk | 0.016 | 0.310 | 0.049 |
| gHusk | 0.164 | 0.188 | 0.072 |
| gPanicle | 0.288 | 0.002 | 0.000 |
| gStraw | 0.220 | 0.056 | 0.044 |
| Ggrain | 0.153 | 0.185 | 0.079 |

#### *ANOVA (Type III tests): Which loading affects Grain Zn?*

| numDF | F-value | p-value | Term |
| --- | --- | --- | --- |
| 1 | 5,289.3791 | 0.0000 | (Intercept) |
| 1 | 5.3433 | 0.0345* | PC1 |
| 1 | 7.4433 | 0.0149* | PC2 |
| 1 | 20.5856 | 0.0003*** | PC3 |
| 1 | 0.5657 | 0.4629 | PmgLFeAUC |
| 3 | 1.3087 | 0.3060 | Climate |

#### *ANOVA (Type III tests): Does climate affect a PC?*

|  | numDF | F-value | p-value |
| --- | --- | --- | --- |
| Intercept | 1 | 0.0000 | 1.0000 |
| PC1 | 3 | 2.0230 | 0.1353 |
| PC2 | 3 | 3.2830 | 0.0366* |
| PC3 | 3 | 1.2913 | 0.2983 |

#### *Model fit statistics:*

| Metric | Value |
| --- | --- |
| AIC | 98.4690 |
| BIC | 105.4223 |
| LogLik | -40.2345 |
| Residual SE | 1.3000 |
| Pseudo-R^2^ | 0.7030 |

### B) **IRON partitioning in high-As soil**

#### *Variance explained by principal components:*

| PC | Std_Dev | Proportion_Variance (%) | Cumulative_Variance |
| --- | --- | --- | --- |
| PC1 | 1.688 | 0.4071 | 0.4071 |
| PC2 | 1.295 | 0.2396 | 0.6467 |
| PC3 | 1.014 | 0.1469 | 0.7936 |
| PC4 | 0.932 | 0.1242 | 0.9178 |
| Other PCs below 10% | | | |

#### *Tissue contributions to principal components (squared loadings, %):*

| Variable | PC1 | PC2 | PC3 | PC4 |
| --- | --- | --- | --- | --- |
| gFeKgStraw | 0.044 | 0.048 | 0.034 | 0.871 |
| mgFeKgPanicle | 0.008 | 0.164 | 0.481 | 0.078 |
| gFeKgHusk | 0.008 | 0.186 | 0.480 | 0.000 |
| gHusk | 0.243 | 0.171 | 0.000 | 0.000 |
| gPanicle | 0.299 | 0.043 | 0.000 | 0.024 |
| gStraw | 0.200 | 0.164 | 0.005 | 0.027 |
| Ggrain | 0.199 | 0.223 | 0.000 | 0.001 |

#### *ANOVA (Type III tests): Which loading affects Grain Zn?*

| numDF | F-value | p-value | Term |
| --- | --- | --- | --- |
| 1 | 2,178.4200 | 0.0000 | (Intercept) |
| 1 | 8.5244 | 0.0100* | PC1 |
| 1 | 4.8984 | 0.0418* | PC2 |
| 1 | 0.0428 | 0.8386 | PmgLFeAUC |
| 3 | 5.1483 | 0.0111* | Climate |

#### *Climate post-hoc comparisons (Tukey-adjusted):*

| contrast | estimate | SE | df | t.ratio | p.value |
| --- | --- | --- | --- | --- | --- |
| Today - eCO_2_ | 2.0577 | 1.1887 | 16 | 1.7311 | 0.3408 |
| Today - eT | -2.2640 | 1.9902 | 16 | -1.1376 | 0.6726 |
| Today - Future | -2.4104 | 1.3122 | 16 | -1.8368 | 0.2931 |
| eCO_2_ - eT | -4.3217 | 1.9232 | 16 | -2.2472 | 0.1528 |
| eCO_2_ - Future | -4.4681 | 1.3955 | 16 | -3.2018 | 0.0257* |
| eT - Future | -0.1464 | 2.3690 | 16 | -0.0618 | 0.9999 |

#### *ANOVA (Type III tests): Does climate affect a PC?*

|  | numDF | F-value | p-value |
| --- | --- | --- | --- |
| Intercept | 1 | 0.0000 | 1.0000 |
| PC1 | 3 | 5.2125 | 0.0059** |
| PC2 | 3 | 4.2669 | 0.0141* |

#### *Model fit statistics:*

| Metric | Value |
| --- | --- |
| AIC | 101.7632 |
| BIC | 107.9439 |
| LogLik | -42.8816 |
| Residual SE | 1.7617 |
| Pseudo-R^2^ | 0.6437 |

### C)  **ZINC partitioning in low-As soil**

#### *Variance explained by principal components:*

| PC | Std_Dev | Proportion_Variance (%) | Cumulative_Variance |
| --- | --- | --- | --- |
| PC1 | 1.721 | 0.4229 | 0.4229 |
| PC2 | 1.274 | 0.2317 | 0.6546 |
| PC3 | 1.094 | 0.1711 | 0.8258 |
| PC4 | 0.864 | 0.1067 | 0.9325 |
| Other PCs below 10% | | | |

#### *Tissue contributions to principal components (squared loadings, %):*

| Variable | PC1 | PC2 | PC3 | PC4 |
| --- | --- | --- | --- | --- |
| mgZnKgStraw | 0.062 | 0.019 | 0.378 | 0.352 |
| mgZnKgPanicle | 0.004 | 0.066 | 0.518 | 0.281 |
| mgZnKgHusk | 0.002 | 0.421 | 0.040 | 0.149 |
| gHusk | 0.237 | 0.130 | 0.002 | 0.093 |
| gPanicle | 0.280 | 0.048 | 0.041 | 0.000 |
| gStraw | 0.182 | 0.187 | 0.015 | 0.065 |
| Ggrain | 0.232 | 0.130 | 0.006 | 0.059 |

#### *ANOVA (Type III tests): Which loading affects Grain Zn?*

| numDF | F-value | p-value | Term |
| --- | --- | --- | --- |
| 1 | 5,706.2564 | 0.0000 | (Intercept) |
| 1 | 15.3468 | 0.0014** | PC1 |
| 1 | 48.4005 | 0.0000*** | PC2 |
| 1 | 23.3489 | 0.0002*** | PC3 |
| 1 | 15.4287 | 0.0013** | PC4 |
| 1 | 2.0878 | 0.1690 | PmgLZnAUC |
| 3 | 5.8767 | 0.0073** | Climate |

#### *Climate post-hoc comparisons (Tukey-adjusted):*

| contrast | estimate | SE | df | t.ratio | p.value |
| --- | --- | --- | --- | --- | --- |
| Today - eCO_2_ | 0.0030 | 1.3890 | 15 | 0.0022 | 1.0000 |
| Today - eT | -5.7355 | 1.6409 | 15 | -3.4953 | 0.0154* |
| Today - Future | -5.5027 | 1.7925 | 15 | -3.0698 | 0.0350* |
| eCO_2_ - eT | -5.7385 | 1.6381 | 15 | -3.5031 | 0.0151* |
| eCO_2_ - Future | -5.5057 | 1.6607 | 15 | -3.3153 | 0.0218* |
| eT - Future | 0.2328 | 1.4624 | 15 | 0.1592 | 0.9985 |

#### *ANOVA (Type III tests): Does climate affect a PC?*

|  | numDF | F-value | p-value |
| --- | --- | --- | --- |
| Intercept | 1 | 0.0000 | 1.0000 |
| PC1 | 3 | 1.8509 | 0.1638 |
| PC2 | 3 | 1.9912 | 0.1411 |
| PC3 | 3 | 4.1184 | 0.0167* |
| PC4 | 3 | 1.0014 | 0.4085 |

#### *Model fit statistics:*

| Metric | Value |
| --- | --- |
| AIC | 104.5314 |
| BIC | 111.6119 |
| LogLik | -42.2657 |
| Residual SE | 2.1418 |
| Pseudo-R^2^ | 0.8907 |

### D) **ZINC partitioning in high-As soil**

#### *Variance explained by principal components:*

| PC | Std_Dev | Proportion_Variance (%) | Cumulative_Variance |
| --- | --- | --- | --- |
| PC1 | 1.759 | 0.4419 | 0.4419 |
| PC2 | 1.222 | 0.2132 | 0.6551 |
| PC3 | 1.111 | 0.1762 | 0.8313 |
| Other PCs below 10% | | | |

#### *Tissue contributions to principal components (squared loadings, %):*

| Variable | PC1 | PC2 | PC3 |
| --- | --- | --- | --- |
| mgZnKgStraw | 0.042 | 0.163 | 0.180 |
| mgZnKgPanicle | 0.001 | 0.065 | 0.512 |
| mgZnKgHusk | 0.108 | 0.242 | 0.000 |
| gHusk | 0.273 | 0.009 | 0.063 |
| gPanicle | 0.215 | 0.170 | 0.048 |
| gStraw | 0.115 | 0.315 | 0.105 |
| Ggrain | 0.247 | 0.036 | 0.092 |

#### *ANOVA (Type III tests): Which loading affects Grain Zn?*

| numDF | F-value | p-value | Term |
| --- | --- | --- | --- |
| 1 | 613.4711 | 0.0000 | (Intercept) |
| 1 | 21.8630 | 0.0003*** | PC1 |
| 1 | 12.8582 | 0.0025** | PC2 |
| 1 | 4.5216 | 0.0494* | PmgLZnAUC |
| 3 | 6.5850 | 0.0042** | Climate |

#### *Climate post-hoc comparisons (Tukey-adjusted):*

| contrast | estimate | SE | df | t.ratio | p.value |
| --- | --- | --- | --- | --- | --- |
| Today - eCO_2_ | 3.9892 | 2.1895 | 16 | 1.8220 | 0.2995 |
| Today - eT | -4.1349 | 2.9166 | 16 | -1.4177 | 0.5069 |
| Today - Future | -4.4036 | 2.2141 | 16 | -1.9889 | 0.2329 |
| eCO_2_ - eT | -8.1242 | 2.5485 | 16 | -3.1878 | 0.0264* |
| eCO_2_ - Future | -8.3928 | 1.9369 | 16 | -4.3332 | 0.0026** |
| eT - Future | -0.2687 | 2.2066 | 16 | -0.1217 | 0.9993 |

#### *ANOVA (Type III tests): Does climate affect a PC?*

|  | numDF | F-value | p-value |
| --- | --- | --- | --- |
| Intercept | 1 | 0.0000 | 1.0000 |
| PC1 | 3 | 6.0907 | 0.0028** |
| PC2 | 3 | 2.2850 | 0.1024 |

#### *Model fit statistics:*

| Metric | Value |
| --- | --- |
| AIC | 109.4918 |
| BIC | 115.6725 |
| LogLik | -46.7459 |
| Residual SE | 2.9841 |
| Pseudo-R^2^ | 0.7867 |

### E)  **CADMIUM partitioning in low-As soil**

#### *Variance explained by principal components:*

| PC | Std_Dev | Proportion_Variance (%) | Cumulative_Variance |
| --- | --- | --- | --- |
| PC1 | 2.077 | 0.6163 | 0.6163 |
| PC2 | 1.236 | 0.2184 | 0.8347 |
| PC3 | 0.762 | 0.0829 | 0.9176 |
| Other PCs below 10% | | | |

#### *Tissue contributions to principal components (squared loadings, %):*

| Variable | PC1 | PC2 | PC3 |
| --- | --- | --- | --- |
| ugCdKgStraw | 0.113 | 0.082 | 0.620 |
| ugCdKgPanicle | 0.101 | 0.282 | 0.003 |
| ugCdKgHusk | 0.070 | 0.264 | 0.222 |
| gHusk | 0.214 | 0.009 | 0.027 |
| gPanicle | 0.181 | 0.075 | 0.070 |
| gStraw | 0.117 | 0.268 | 0.048 |
| Ggrain | 0.205 | 0.020 | 0.009 |

#### *ANOVA (Type III tests): Which loading affects Grain Zn?*

| numDF | F-value | p-value | Term |
| --- | --- | --- | --- |
| 1 | 332.0215 | 0.0000 | (Intercept) |
| 1 | 49.9026 | 0.0002*** | PC1 |
| 1 | 14.4918 | 0.0067** | PC2 |
| 1 | 0.0337 | 0.8596 | PmgLCdAUC |
| 3 | 3.0643 | 0.1006 | Climate |

#### *ANOVA (Type III tests): Does climate affect a PC?*

|  | numDF | F-value | p-value |
| --- | --- | --- | --- |
| Intercept | 1 | 0.0000 | 1.0000 |
| PC1 | 3 | 11.637 | 0.001** |
| PC2 | 3 | 2.2559 | 0.1389 |

#### *Model fit statistics:*

| Metric | Value |
| --- | --- |
| AIC | 45.1736 |
| BIC | 44.7408 |
| LogLik | -14.5868 |
| Residual SE | 1.8049 |
| Pseudo-R^2^ | 0.9132 |

### F)  **CADMIUM partitioning in high-As soil**

#### *Variance explained by principal components:*

| PC | Std_Dev | Proportion_Variance (%) | Cumulative_Variance |
| --- | --- | --- | --- |
| PC1 | 1.994 | 0.5683 | 0.5683 |
| PC2 | 1.244 | 0.2212 | 0.7895 |
| PC3 | 0.974 | 0.1357 | 0.9251 |
| Other PCs below 10% | | | |

#### Tissue contributions to principal components (squared loadings, %):

| Variable | PC1 | PC2 | PC3 |
| --- | --- | --- | --- |
| ugCdKgStraw | 0.064 | 0.353 | 0.084 |
| ugCdKgPanicle | 0.039 | 0.007 | 0.839 |
| ugCdKgHusk | 0.081 | 0.285 | 0.029 |
| gHusk | 0.188 | 0.149 | 0.000 |
| gPanicle | 0.235 | 0.006 | 0.028 |
| gStraw | 0.219 | 0.018 | 0.018 |
| Ggrain | 0.174 | 0.182 | 0.002 |

#### *ANOVA (Type III tests): Which loading affects Grain Zn?*

| numDF | F-value | p-value | Term |
| --- | --- | --- | --- |
| 1 | 107.2971 | 0.0000 | (Intercept) |
| 1 | 0.5987 | 0.4589 | PC1 |
| 1 | 20.6180 | 0.0014** | PC2 |
| 1 | 3.0711 | 0.1136 | PmgLCdAUC |
| 3 | 1.2765 | 0.3401 | Climate |

#### *ANOVA (Type III tests): Does climate affect a PC?*

|  | numDF | F-value | p-value |
| --- | --- | --- | --- |
| Intercept | 1 | 0.0000 | 1.0000 |
| PC2 | 3 | 7.3044 | 0.0048** |

#### *Model fit statistics:*

| Metric | Value |
| --- | --- |
| AIC | 60.6594 |
| BIC | 62.2372 |
| LogLik | -22.3297 |
| Residual SE | 2.4601 |
| Pseudo-R^2^ | 0.7575 |

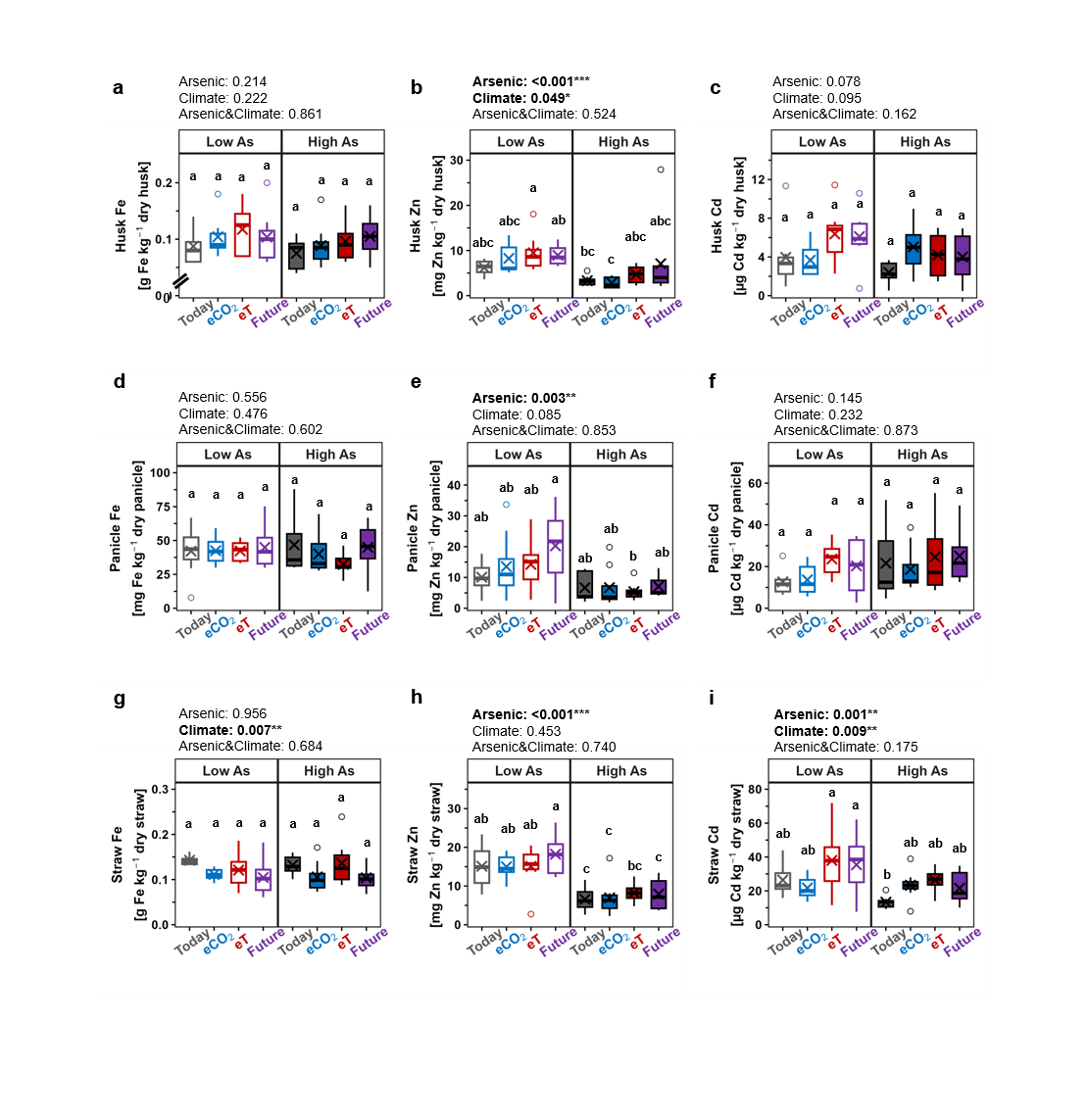

**Fig. S1** Husk, panicle, and straw elemental concentrations in *Oryza sativa* L. cv. M206 grown under varying climatic and soil arsenic conditions. Husk concentrations of (a) iron, (b) zinc, and (c) cadmium; panicle concentrations of (d) iron, (e) zinc, and (f) cadmium; and straw concentrations of (g) iron, (h) zinc, and (i) cadmium are provided. Rice was subjected to today’s (33ºC, 415 ppmv CO_2_; grey), elevated CO_2_ (33ºC, 850 ppmv CO_2_; blue), elevated temperature (38ºC, 415 ppmv CO_2_; red), and future (38ºC, 850 ppmv CO_2_; purple) climatic conditions with low (no fill, 7.3 mg As kg^−1^_dry soil_) and high (filled, amended to 24.5 mg As kg^−1^_dry soil_) soil arsenic levels. Each treatment included eight biological replicates. Boxplots show the median, interquartile range (IQR), and whiskers extending to the most extreme values within 1.5 × IQR; observations outside this range are plotted as outliers (circles). Statistical analyses of climate, arsenic, and their interaction on elemental grain concentrations were performed using ANOVA at a 95% confidence level using the results of a generalized least squares test. Results are shown above each plot, with significant findings highlighted in bold. Post-hoc comparisons using the Emmeans test are indicated by different lowercase letters, where letters denote statistically significant differences (p<0.05).

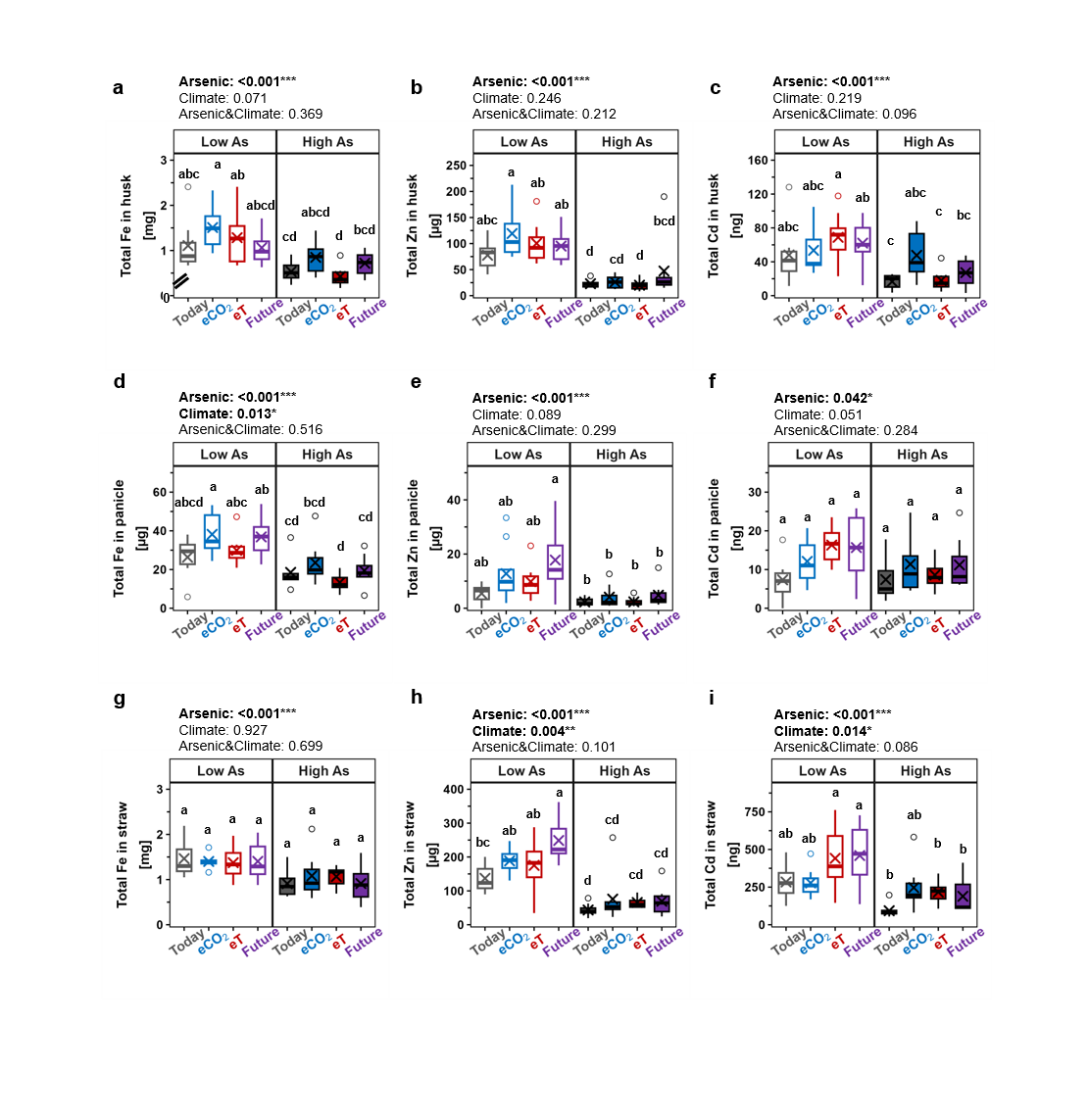

**Fig. S2** Total husk, panicle, and straw elemental contents in *Oryza sativa* L. cv. M206 grown under varying climatic and soil arsenic conditions. Total elemental contents in husks for (a) iron, (b) zinc, and (c) cadmium; total elemental contents in panicles for (d) iron, (e) zinc, and (f) cadmium; and total elemental contents in straw for (g) iron, (h) zinc, and (i) cadmium are provided. . Rice was subjected to today’s (33ºC, 415 ppmv CO_2_; grey), elevated CO_2_ (33ºC, 850 ppmv CO_2_; blue), elevated temperature (38ºC, 415 ppmv CO_2_; red), and future (38ºC, 850 ppmv CO_2_; purple) climatic conditions with low (no fill, 7.3 mg As kg^−1^_dry soil_) and high (filled, amended to 24.5 mg As kg^−1^_dry soil_) soil arsenic levels.. Each treatment included eight biological replicates. Boxplots show the median, interquartile range (IQR), and whiskers extending to the most extreme values within 1.5 × IQR; observations outside this range are plotted as outliers (circles). Statistical analyses of climate, arsenic, and their interaction on elemental grain contents were performed using ANOVA at a 95% confidence level using the results of a generalized least squares test. Results are shown above each plot, with significant findings highlighted in bold. Post-hoc comparisons using the Emmeans test are indicated by different lowercase letters, where letters denote statistically significant differences (p<0.05).

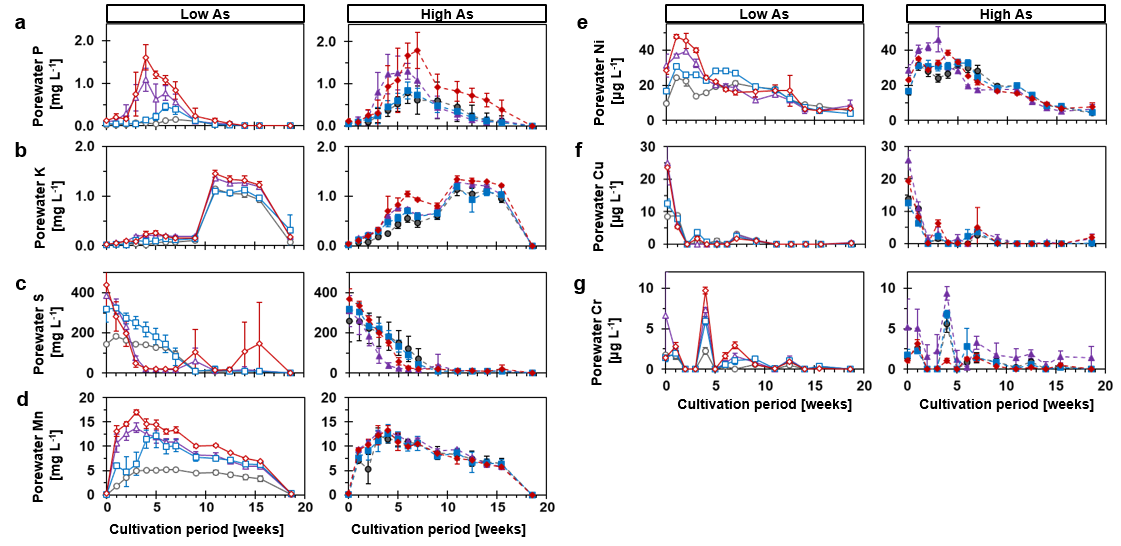

**Fig. S3** Time-resolved dissolved elements in rhizosphere porewater of Oryza sativa L. cv. M206 grown under varying climatic and soil arsenic conditions. Total porewater (a) phosphorous, (b) potassium, (c) sulfur, (d) manganese, (e) nickel, (f) copper, and (g) chromium extracted from the rhizosphere of rice plants at 10-cm depth are presented. . Rice was subjected to today’s (33ºC, 415 ppmv CO_2_; grey), elevated CO_2_ (33ºC, 850 ppmv CO_2_; blue), elevated temperature (38ºC, 415 ppmv CO_2_; red), and future (38ºC, 850 ppmv CO_2_; purple) climatic conditions with low (no fill, 7.3 mg As kg^−1^_dry soil_) and high (filled, amended to 24.5 mg As kg^−1^_dry soil_) soil arsenic levels. At each sampling point, the mean ± standard deviation of six biological replicates was calculated.

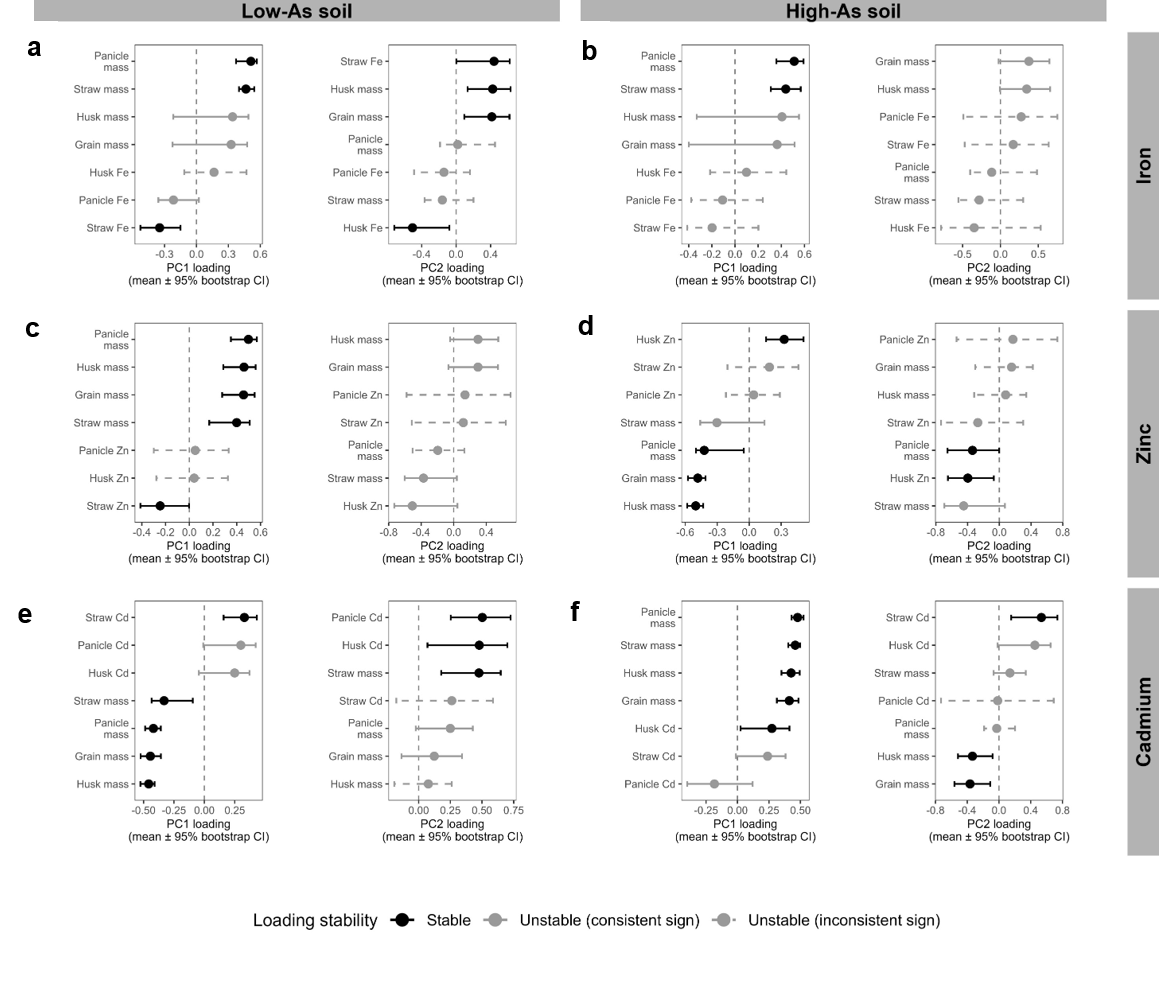

**Fig. S4** Bootstrap stability of PCA loadings across element partitioning systems in rice plants grown on arsenic-bearing soils. PC1 and PC2 bootstrap stabilities of (a,b) iron, (c,d) zinc, and (e,f) cadmium in rice grown on low- (left panels, 7.3 mg As kg^−1^_dry soil_) and high- (right panels, 24.5 mg As kg^−1^_dry soil_) arsenic soils. Bootstrap resampling (n = 1000) was used to assess the robustness of tissue loadings on principal components describing element partitioning among plant tissues. Points represent mean loadings across bootstrap iterations, with horizontal error bars indicating 95% bootstrap confidence intervals. Loadings whose confidence intervals do not overlap zero are classified as stable (black). Loadings with confidence intervals crossing zero are shown in grey and further classified based on directional consistency across bootstrap samples: solid lines indicate high sign consistency (≥90% agreement in loading sign), whereas dashed lines indicate low sign consistency (<90%), indicating frequent sign switching. This classification separates uncertainty in loading magnitude from instability in loading direction, enabling comparison of PCA robustness across elements and soil arsenic conditions.

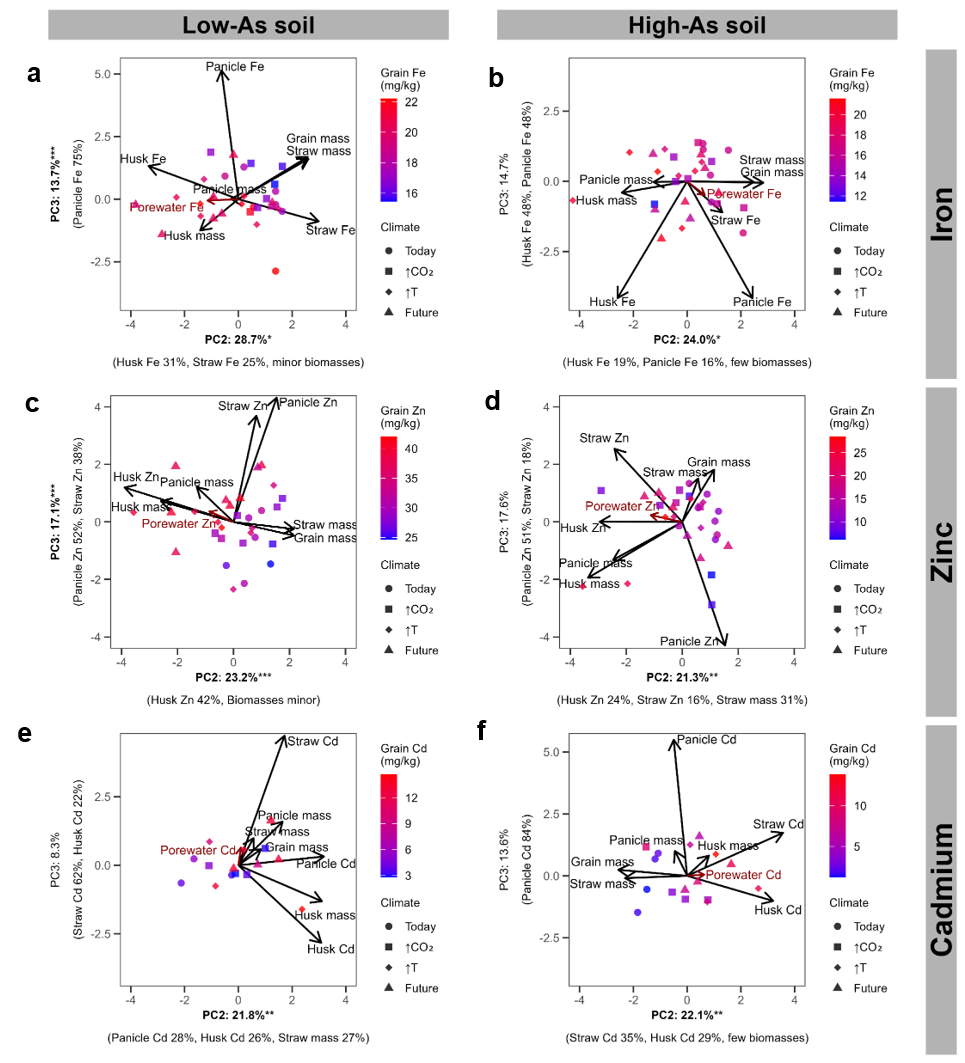

**Fig. S5** Elemental partitioning structure PC2-PC3 of rice grown on arsenic-bearing paddy soils under climate change conditions. Principal component analyses (PCA) of (a,b) iron, (c,d) zinc, and (e,f) cadmium partitioning among rice tissues grown in low-As (left panels) and high-As (right panels) soils under today’s (circle), elevated CO₂ (↑CO₂, squares), elevated temperature (↑T, diamond), and combined future (triangle) climate conditions. PCA was performed on scaled tissue element concentrations (straw, panicle and husk) together with biomass allocation traits (straw, panicle, husk and grain mass) to characterize whole-plant elemental distribution independent of absolute magnitude. Points represent individual experimental replicates, colored by grain element concentration and shaped by climate condition. Arrows denote variable loadings in PC2-PC3 space indicating the direction and strength of tissue contributions to multivariate partitioning axes. The red vector represents porewater elemental exposure projected onto PCA space, illustrating the alignment between environmental availability and plant partitioning strategies. Axis labels indicate the percentage of explained variance and dominant tissue contributors. Bold axis labels with significance symbols denote PCs that significantly explained variation in grain element concentration in generalized least-squares models (*p < 0.05, **p < 0.01, ***p < 0.001). Together, the first two PCs capture dominant trade-offs between biomass allocation and tissue elemental enrichment across As and climate conditions.

**Note S1** Supplementary discussion of porewater data.

Though porewater concentrations of Fe, Zn, and Cd did not drive grain concentrations (Fig. 3), they provided insight into the factors controlling their phytoavailability under soil As contamination and future climate. In waterlogged paddy soils, microbial oxygen consumption drives the use of alternative electron acceptors under anoxic conditions, e.g. Mn and Fe (Zobrist et al. 2000; Kocar et al. 2006; Blume et al. 2016; Sharma et al. 2021). This promotes reductive dissolution of minerals, which was evident in the sharp increase in porewater Mn (weeks 1-3; Fig. S3d) followed by increasing porewater Fe (Fig. 2a). This is further evidenced by the co-occurring increase in porewater P (Fig. S3a), an element which is known to be closely associated with Fe minerals and is released upon their reductive dissolution (Blume et al. 2016). When reactive metal oxides were depleted (~week 5 in future and ~week 15 in today’s climate), adsorption to soil solid phases, precipitation as FeS, and plant and microbial uptake led to gradually decreasing concentrations of porewater Fe and Mn (Fig.s 2a, S3d).

Porewater Zn followed a similar trend as S, likely indicating its solubility was controlled by equilibrium between dissolution and precipitation in the common form in paddy soils, ZnS. Chemical weathering of silicates can release cations such as K^+^ (Blume et al. 2016), thus the large peak in potassium after week 9 is likely linked to weathering of clay minerals. Cd followed a similar trend as Zn, likely due to release of Cd from weathering of clay minerals. Porewater investigation indicated distinct solid phases controlling phytoavailability of Fe, Zn, and Cd in the paddy soil, with Fe solubility linked to reductive dissolution and Zn and Cd solubility linked to (de)sorption to(from) clay minerals and dissolution/precipitation of metal sulfides (ZnS, CdS).
